# Integrative molecular roadmap for direct conversion of fibroblasts into myocytes and myogenic progenitor cells

**DOI:** 10.1101/2021.08.20.457151

**Authors:** Inseon Kim, Adhideb Ghosh, Nicola Bundschuh, Laura Hinte, Ferdinand von Meyenn, Ori Bar-Nur

## Abstract

Transient MyoD overexpression in concert with small molecules treatment reprograms mouse fibroblasts into induced myogenic progenitor cells (iMPCs). However, the molecular landscape and mechanisms orchestrating this cellular conversion remain unknown. Here, we undertook an integrative multi-omics approach to delineate the process of iMPC reprogramming in comparison to myogenic transdifferentiation mediated solely by MyoD. Utilizing transcriptomics, proteomics and genome-wide chromatin accessibility assays, we unravel distinct molecular trajectories which govern the two processes. Notably, iMPC reprogramming is characterized by gradual upregulation of stem and progenitor cell markers, unique signaling pathways, chromatin remodelers and cell cycle regulators which manifest via rewiring of the chromatin in core myogenic promoters. Furthermore, we determine that only iMPC reprogramming is mediated by Notch pathway activation, which is indispensable for iMPC formation and self-renewal. Collectively, this study charts divergent molecular blueprints for myogenic transdifferentiation or reprogramming and underpins the heightened capacity of iMPCs in capturing myogenesis *ex vivo.*

## Introduction

Skeletal muscle is a soft tissue which governs voluntary movement and accounts for 30-40% of the normal human body mass. This tissue is predominately composed of multinucleated muscle fibers that contract to generate locomotion and a variety of mononucleated resident cells that maintain tissue homeostasis^1^. Satellite cells are resident skeletal muscle stem cells that can regenerate muscle fibers upon injury or disease and are characterized by high expression of the transcription factor paired box protein 7 (Pax7)^2–4^. These cells reside in a unique anatomical location between the myofiber cell membrane and basal lamina and are quiescent during homeostasis, undergoing activation to repair tissue damage following muscle insult^2,3^. During this repair process, activated satellite cells either divide symmetrically to increase the pool of proliferative satellite cells that eventually return to quiescence, or divide asymmetrically into myoblasts and fusion-competent myocytes that merge and repair damaged myofibers^5^. Satellite cell activation is a stepwise process which commences by upregulation of myogenic regulatory transcription factors including Myf5 and MyoD in myoblasts, Myogenin (Myog) and Myf6/MRF4 in myocytes and Myosin heavy chain (MyHC) isoforms in multinucleated myofibers^2,3^. Following isolation from skeletal muscles and *in vitro* propagation, satellite cells form a population of proliferative myoblasts which upregulate MyoD and rapidly lose molecular attributes indicative of an *in vivo* activated satellite cell state^6^. This loss of satellite cell attributes following *in vitro* expansion renders myoblasts cumbersome for regenerative medicine purposes and highlights the necessity to seek alternative methods to culture myogenic stem and progenitor cells^7,8^.

Direct lineage reprogramming denotes the conversion of one cell type into another. It is typically induced by forced overexpression of cell-type-specific transcription factors or small molecule treatment^9^. Manipulation of cell identity via this approach was first demonstrated in a milestone study which determined that overexpression of the transcription factor MyoD transdifferentiates fibroblasts into skeletal muscle cells^10^. Since this seminal study, several works reported on transdifferentiation of somatic cells into various cell types including neurons, cardiomyocytes and hepatocytes^11–14^. Moreover, further works have reported on direct reprogramming of somatic cells into tissue-specific multipotent stem and progenitor cells^15–20^. One study recently reported on a method to directly reprogram mouse fibroblasts into “induced myogenic progenitor cells” (iMPCs) by transient MyoD overexpression in conjunction with three small molecules; the adenylate cyclase activator Forskolin (F), the TGF-β receptor inhibitor RepSox (R) and the GSK3-β inhibitor CHIR99021 (C) (abbreviated as F/R/C)^20^. Reprogramming into iMPCs is markedly different from conventional transdifferentiation into myogenic cells solely by MyoD, which typically only generates multinucleated myotubes^20^. In contrast, MyoD overexpression in concert with F/R/C treatment gives rise to heterogeneous and expandable myogenic cultures consisting of skeletal muscle progenitor cells that express Pax7 and Myf5, in addition to highly contractile myofiber netowork^20^.

The divergent lineage conversion trajectories between MyoD and MyoD+F/R/C treatment raises the question how small molecules administration endows a myogenic stem cell fate on fibroblasts in comparison to postmitotic myotubes by MyoD-mediated transdifferentiation, which has been extensively studied^21–23^. Furthermore, it remains unknown how molecularly akin are Pax7^+^ iMPCs to primary Pax7^+^ myoblasts. To address these questions, here we set out to delineate the molecular landscape of iMPCs utilizing integrative multi-omics approaches and further dissect the molecular trajectory guiding fibroblast conversion into iMPCs in comparison to transdifferentiation solely by MyoD. Collectively, we establish a body of knowledge in respect to the genes, proteins and pathways that govern direct lineage conversion into myogenic progenitor cells, and further establish their molecular disparity from primary myoblasts. Our results suggest that iMPCs capture a *bona fide* skeletal muscle differentiation program *in vitro*, thus establishing them as an exceptional method to culture and propagate skeletal muscle stem and progenitor cells.

## Results

### An inducible reprogramming system to study fibroblast conversion into myogenic cells

We commenced our investigation by establishing an inducible cellular conversion system in fibroblasts that enables comparison of reprogramming via MyoD+F/R/C to transdifferentiation solely by MyoD. To this end, we derived mouse embryonic fibroblasts (MEFs) from a transgenic mouse strain that carries a *Pax7-nuclear GFP* (*Pax7-nGFP*) reporter, which allows prospective skeletal muscle stem cell purification using fluorescence-activated cell sorting (FACS)^24^. To initiate controlled MyoD overexpression in *Pax7-nGFP* MEFs, we engineered a novel doxycycline (dox)-inducible Tet-On lentiviral dual plasmid system harboring a MyoD coding sequence under a TRE3G promoter (*tetO-MyoD/PGK-Puromycin*) and a Tet3G activator under the control of a constitutive EF1-α promoter (*EF1*α*-rtTA3/PGK-Neomycin)* (Fig. 1A). Each respective plasmid also contained an antibiotic resistance gene, enabling selection of reprogrammable MEFs (Rep-MEFs) that carry both lentiviral constructs (Fig. 1A). Since MEFs might contain multiple cell types, we FACS-purified fibroblasts from MEF cultures using Thy1, a fibroblast-specific marker (Supplementary Fig. 1A)^25^. We then subjected Thy1^+^ *Pax7-nGFP* Rep-MEFs to either dox (MyoD) or dox+F/R/C (MyoD+F/R/C) treatment. Following 1-3 days of MyoD overexpression with or without F/R/C treatment, we documented the formation of multinucleated myotubes, however from days 6-8, Rep-MEFs treated with MyoD+F/R/C formed a highly contractile network of myofibers in conjunction with appearance of small mononucleated cells which proliferated robustly (Fig. 1B). To assess the expression of myogenic markers upon MyoD or MyoD+F/R/C treatment, we performed quantitative Real-Time PCR (qRT-PCR) at day 10 of reprogramming and documented robust upregulation of the myogenic stem cell markers *Pax7* and *Myf5* in MyoD+F/R/C treatment, whereas the differentiation marker *Myog* was upregulated in both conditions (Fig. 1C). Significant downregulation of the fibroblast marker *Col1a1* was observed only in MyoD+F/R/C condition (Supplementary Fig. 1B).

**Fig. 1:**
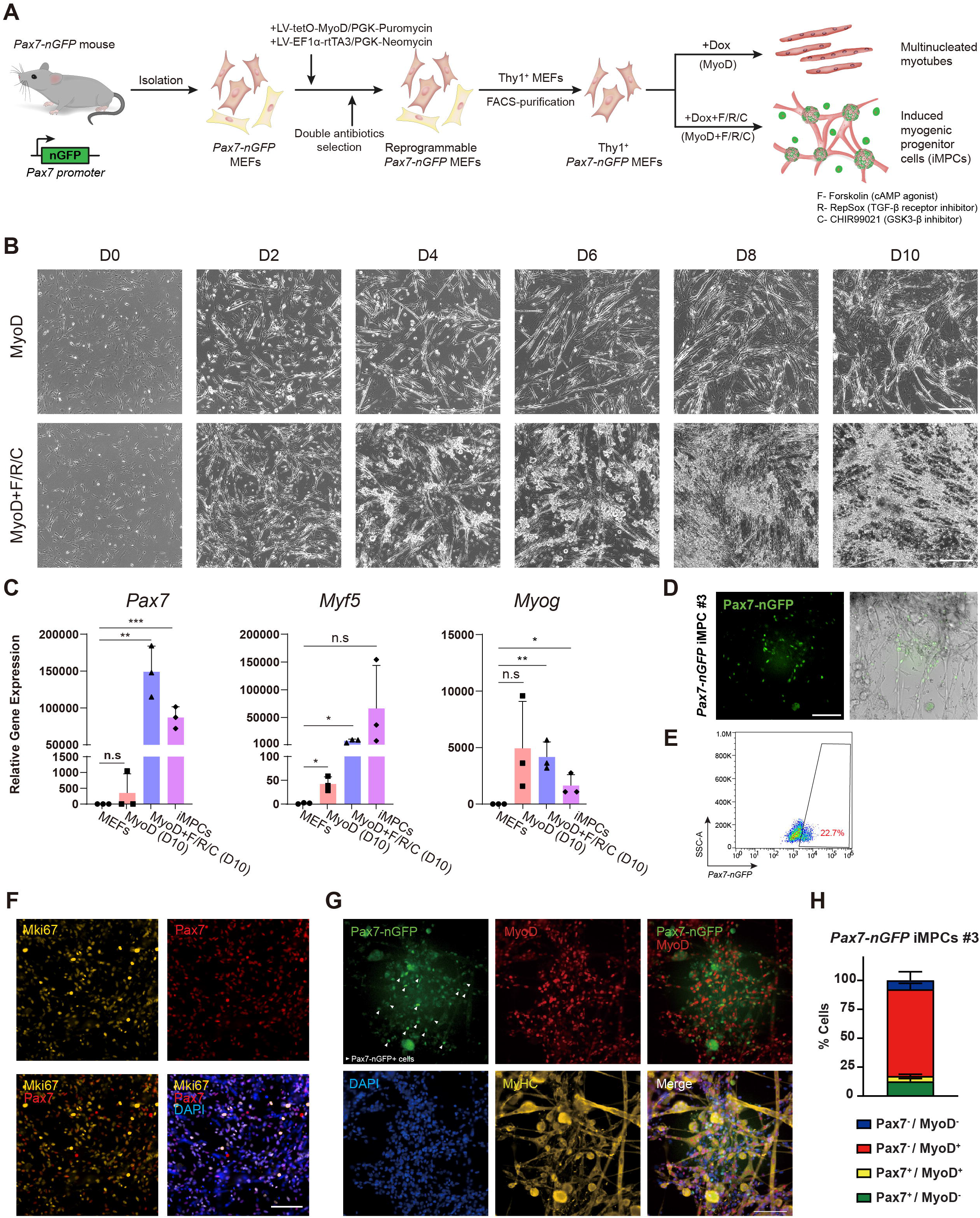
A reprogrammable system to study fibroblasts conversion into myogenic cells. **(A)** A schematic of experimental design denoting the conversion of Thy1^+^ *Pax7-nGFP* Rep-MEFs into multinucleated myotubes solely by MyoD overexpression or iMPCs by MyoD+F/R/C treatment. MEFs, mouse embryonic fibroblasts; Dox, Doxycycline. **(B)** Representative bright-field images of Thy1^+^ Rep-MEFs subjected to MyoD or MyoD+F/R/C treatment for the indicated days. Scale bar, 400 μm. **(C)** qRT-PCR for canonical myogenic genes. Data is shown as means□±□S.D. N = 3 cell lines per group. Statistical significance was determined by a two-tailed unpaired *t*-test (*p<0.05, **p<0.01, ***p<0.001, n.s=non-significant). **(D)** Representative images of a stable *Pax7-nGFP* iMPC clone at passage 1. Scale bar, 100 μm. **(E)** Flow cytometry analysis of a *Pax7-nGFP* iMPC clone. **(F)** Representative immunofluorescence images of *Pax7-nGFP* iMPCs immunostained for Mki67 and Pax7. Nuclei were counterstained with 4′,6-diamidino-2-phenylindole (DAPI). Scale bar, 100 μm. **(G)** Representative immunofluorescence images of *Pax7-nGFP* iMPCs immunostained for MyoD and MyHC. Nuclei were counterstained with DAPI. White arrowheads point to mononucleated *Pax7-nGFP^+^* cells that are negative for MyoD. Scale bar, 100 μm. **(H)** Quantification of (f). N = 4 field images that were taken from the respective iMPC line.

Next, to generate stable dox-independent iMPC clones we reprogrammed Rep-MEFs using MyoD+F/R/C treatment for 10 days and then withdrew dox and expanded the cells in medium containing F/R/C for several passages, thus establishing dox-independent iMPC clones which proliferated robustly and expressed an array of myogenic progenitor and differentiation genes (Fig. 1C, D). Furthermore, these cultures contained a network of contractile myofibers as well as mononucleated Pax7-nGFP^+^ cells, ranging between 3.6-22.7%, that were also positive for the cell proliferation marker Mki67 (Fig. 1D, E, F Supplementary Fig. 1C, D). Immunostaining for MyoD and MyHC revealed the presence of Pax7-nGFP^+^/MyoD^−^/MyHC^−^ mononucleated cells, suggesting that iMPCs contain more immature myogenic stem / progenitor cells as previously reported (Fig. 1G)^20^. Quantification of the cell subsets in an established iMPC clone revealed the presence of Pax7^+^/MyoD^−^ (12.6 ± 2.5%), Pax7^+^/MyoD^+^ (4.7 ± 1.5%), Pax7^−^/MyoD^+^ (74.9 ± 5.3%) and Pax7^−^/MyoD^−^ (7.9 ± 7.4%) cells, highlighting the heterogeneous cell composition of iMPCs (Fig. 1H). Collectively, we successfully established a novel reprogramming system to interrogate transdifferentiation by MyoD or reprogramming by MyoD+F/R/C. Furthermore, we demonstrate that subjecting fibroblasts to MyoD+F/R/C treatment forms highly heterogeneous iMPC cultures consisting of mononucleated Pax7-nGFP^+^ cells that can propagate without ectopic MyoD overexpression.

### Transcriptional dynamics during fibroblast conversion into skeletal muscle cells

To gain molecular insights into the transcriptional changes which occur during iMPC formation, we opted to perform bulk RNA-Sequencing (RNA-Seq) and dissect transcriptional dynamics following MyoD or MyoD+F/R/C treatment of fibroblasts. To this end, we subjected Thy1^+^ Rep-MEFs to MyoD or MyoD+F/R/C treatment, followed by RNA-Seq at days 2, 4, 6, 8, and 10 of the conversion process (Fig. 2A). As positive controls, we used iMPC clones and satellite cell-derived *Pax7-nGFP* primary myoblasts. Principal Component Analysis (PCA) and dendrogram clustering separated the MyoD and MyoD+F/R/C treated cells into two distinct groups, which were both transcriptionally divergent from parental MEFs (Fig. 2B, Supplementary Fig. 2A). In the PCA, a stepwise temporal trajectory was documented during MyoD+F/R/C reprogramming course, as cells gradually clustered further away from parental MEFs and became more akin to stable iMPCs and primary myoblasts (Fig. 2B). In contrast, MEFs subjected only to MyoD overexpression exhibited a haphazard temporal trajectory with no clear separation between time points (Fig. 2B). We next documented the number of differentially expressed genes (DEGs) during MyoD or MyoD+F/R/C conditions. This analysis revealed a prominent transcriptional wave of upregulated genes in both conditions at day 2 and 4, followed by reduced transcriptional activity during days 4-6, and an additional transcriptional wave only in MyoD+F/R/C treated MEFs at day 8 (Fig. 2C). We then investigated individual gene expression dynamics during the reprogramming course and noted upregulation of the differentiation markers *Myog*, *Tnni1, Actn2* and *Myh2* in both MyoD and MyoD+F/R/C conditions (Fig. 2D, Supplementary Fig. 2B). However, prominent expression of other key myogenic progenitor and differentiation markers including *Myf5*, *Myf6, Six1, Six4, Myh1* and *Myh4* was significantly upregulated only in MyoD+F/R/C condition (Fig. 2D, Supplementary Fig.2B). Notably, significant upregulation of immature myogenic stem and progenitor markers including *Pax7*, *Dmrt2* and *Sox8* was only observed in MyoD+F/R/C condition starting at day 8 of the reprogramming process (Fig. 2D, Supplementary Fig. 2C). Additionally, pronounced downregulation of fibroblast-specific genes such as *Fbln2*, *Col1a1* and *Col5a1* was most notably observed in the presence of F/R/C treatment, whereas *Thy1* expression was significantly downregulated in both conditions (Fig. 2D, Supplementary Fig. 2B, C).

**Fig. 2:**
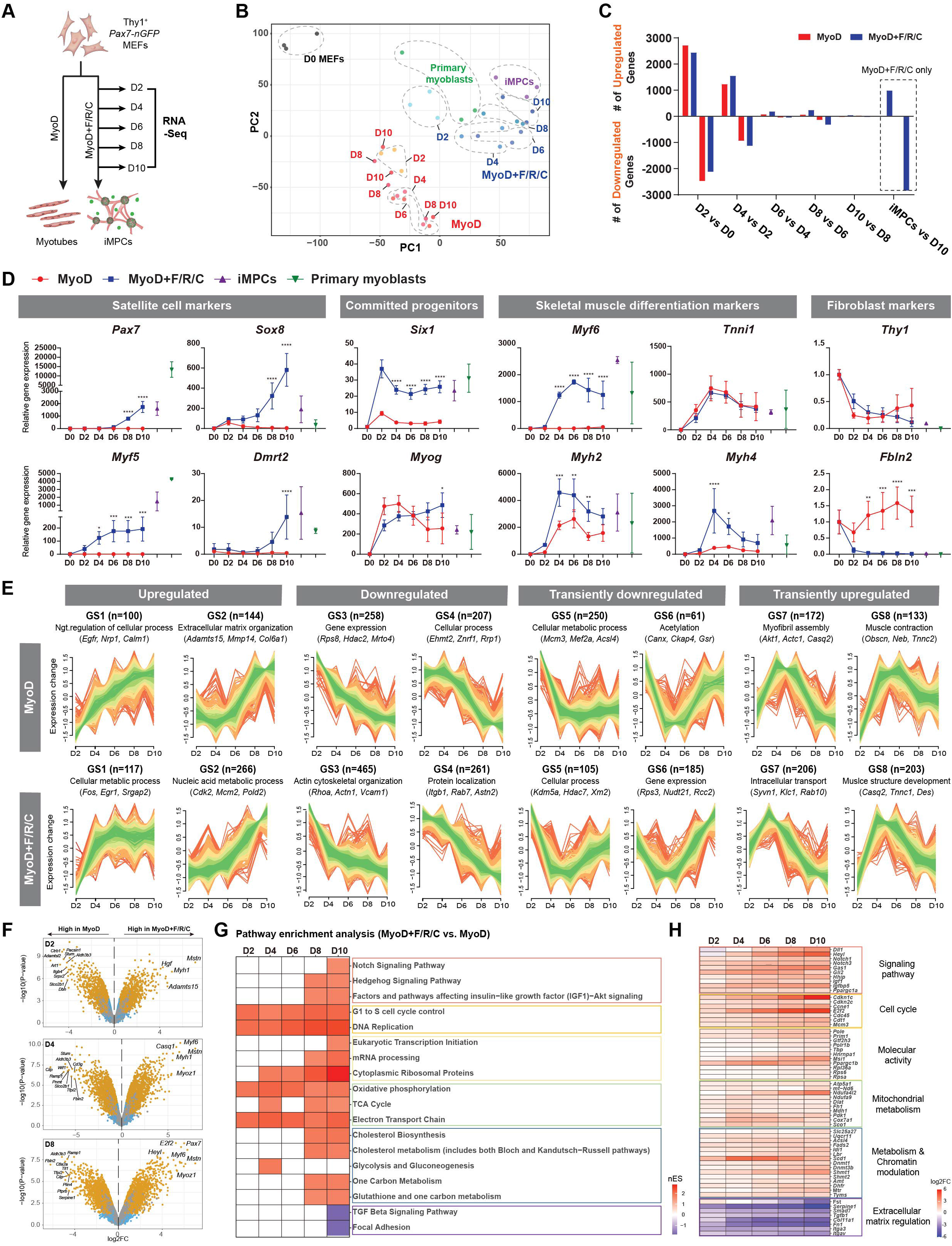
Global transcriptome dynamics during myogenic reprogramming and transdifferentiation. **(A)** An experimental design for the bulk RNA-Seq analysis. **(B)** PCA of global RNA-Seq data using all gene read counts. N = 3 cell lines per group. **(C)** Graph showing the numbers of DEGs between each indicated comparison. DEGs were calculated using |log2FC|>0.5 (p-value<0.01). DEGs, differentially expressed genes. **(D)** Gene expression dynamics for the indicated markers based on bulk RNA-Seq data during MyoD or MyoD+F/R/C conversion. Established iMPCs and primary myoblasts served as positive controls. Relative gene expression was calculated by normalizing the Reads per Kilobase Million (RPKM) values of each sample to that of parental MEFs (D0 MEFs). The data is shown as meansL±LS.D. N = 3 cell lines per group. Statistical significance was determined by two-way ANOVA between conditions at each time point (*p<0.05, **p<0.01, ***p<0.001, ****p<0.0001). **(E)** Fuzzy clustering based on bulk RNA-Seq data for gene expression dynamics across the indicated days in MEFs subjected to MyoD or MyoD+F/R/C treatment. Expression changes are represented as log2FC for each time point vs. parental MEFs. N = 3 cell lines per each group. The number of genes is shown together with the respective GO term annotation and representative genes in each gene set. GO term and gene annotations were defined using the STRING database. **(F)** Volcano plots showing DEGs between MyoD and MyoD+F/R/C conditions at the indicated days. Significant DEGs (|log2FC|>0.5, p-value<0.05) are shown as yellow dots. N = 3 cell lines per group. **(G)** A heatmap of enriched pathways in MyoD+F/R/C vs. MyoD conditions at the indicated time points. Normalized enrichment score (nES) obtained via Gene Set Enrichment Analysis (GSEA) using WikiPathways database is displayed. N = 3 cell lines per group. **(H)** A heatmap of relative gene expression based on bulk RNA-Seq data for each of the indicated genes at the respective time points. Each gene group is associated with the respective pathway shown in (E). Log2FC was calculated for MyoD+F/R/C vs. MyoD condition. N = 3 cell lines per group.

We next analyzed broad transcriptional trends in the two cellular conversions via fuzzy clustering. In total, 8 gene subsets that exhibited similar expression patterns during MyoD or MyoD+F/R/C conditions were defined (Fig. 2E). The first group involved two gene sets (GS1, GS2) which were upregulated in both conditions, albeit for the MyoD condition these genes annotated with *cellular processes*, whereas *metabolic processes* were defined for the MyoD+F/R/C condition (Fig. 2E). Specifically, 266 genes in GS2 of MyoD+F/R/C treatment exhibited a gradual gene upregulation pattern over time and contained the subsets annotated with *Regulation of cell cycle (Cdk2*, *Cdkn1c*, *Smc3)* and *Striated muscle tissue development* (*Pax7*, *Heyl*, *Tbx3*) (Supplementary Fig. 2D, E). The second gene sets (GS3, GS4) involved downregulated genes in both conditions and in particular GS3 exhibited downregulation of the largest subsets of genes over time (Fig. 2E). Lastly, gene subsets showing transient downregulation (GS5, GS6) or upregulation (GS7, GS8) patterns were defined (Fig. 2E).

The transcriptome analysis revealed prominent gene expression differences between MyoD and MyoD+F/R/C treatment. Given this observation, we next carried out differential gene expression analysis between the two cell conversions to uncover key DEGs. The top 30 upregulated genes in MyoD or MyoD+F/R/C conditions (day 10) vs. MEFs were associated with skeletal muscle differentiation markers including *Trim72*, *Tnnt1*, *Tnni1*, *Tnnc2*, *Neb*, *Myh7*, *Myh8* and *Actn2* (Supplementary Fig. 3A). However, the top 30 upregulated genes in MyoD+F/R/C vs. MyoD conditions at day 10 were mainly myogenic stem and progenitor cell markers including *Pax7*, *Sox8*, *Msc*, *Lgr5*, *Fgfr4*, *Dbx1* and *Heyl* (Supplementary Fig. 3A). Given the elevated expression of myogenic stem cell markers in MyoD+F/R/C in comparison to MyoD at day 10, we next performed sequential analysis of DEGs in MyoD+F/R/C vs. MyoD conditions. The most upregulated genes at days 2, 4, and 6 in MyoD+F/R/C were predominantly skeletal muscle differentiation markers including *Mstn*, *Myoz1*, *Myh1*, *Myf6* and *Casq1* (Fig. 2F Supplementary Fig. 3B, 3C). The most differentially upregulated genes at a late reprogramming stage of MyoD+/F/R/C conditions (days 8 and 10) consisted of stem cell markers including *Pax7*, *Sox8*, *Dbx1* and the cell cycle regulator *Cdkn1c*, in addition to the differentiation markers *Myf6*, *Mstn* and *Myoz1* (Fig. 2F, Supplementary Fig. 3B). To further validate the expression of muscle stem cell genes during MyoD+F/R/C reprogramming, we performed a meta comparison of the transcriptomes of *Pax7-nGFP* primary myoblasts with MEFs subjected to MyoD or MyoD+F/R/C conditions. Of note, we documented that satellite cell-associated genes are only shared between primary myoblasts and MyoD+F/R/C treated cells during days 6-10, whereas several differentiation markers were detected in all conditions for all inspected time points (Supplementary Fig. 3D). To acquire additional insights into the signaling pathways which are enriched during MyoD+F/R/C treatment, we performed pathway enrichment analysis of DEGs between MyoD+F/R/C vs. MyoD conditions using gene set enrichment analysis (GSEA) for each reprogramming time point. Pathways enriched in MyoD+F/R/C condition across all time points were associated with “*Cell cycle*” and “*Mitochondrial metabolism*”, whereas “*Metabolism/chromatin modulation*” and “*Notch and Hedgehog signaling pathways*” were enriched in MyoD+F/R/C condition only at days 8 and 10 (Fig. 2G, H).

In summary, using bulk RNA-Seq we delineated transcriptional changes during cellular reprogramming manifested by MyoD or MyoD+F/R/C treatment and demonstrated divergent transcriptional dynamics. Solely overexpressing MyoD in fibroblasts induces fast and direct formation of multinucleated myotubes that express only a partial cohort of myogenic differentiated genes. In contrast, MyoD+F/R/C treatment entails a stepwise reprogramming process which commences with an early transcriptional wave that is characterized by upregulation of a plethora of myogenic differentiation genes prior to a second transcriptional wave that is characterized by expression of myogenic stem and progenitor cell markers. Furthermore, direct transcriptional comparison between MyoD+F/R/C and MyoD conditions at defined time points revealed canonical stem cell and differentiation markers in addition to signaling pathways that are unique to iMPC reprogramming.

### Proteome dynamics during direct conversion of fibroblasts into myogenic cells

The RNA-Seq analysis revealed pronounced transcriptional differences between MEFs subjected to MyoD or MyoD+F/R/C conditions. As such, we next wished to assess whether these differences might be reflected at the protein level. To this end, we performed Liquid chromatography-mass spectrometry (LC-MS) analysis of multiple MEFs, iMPC clones as well as MEFs subjected to MyoD or MyoD+F/R/C conditions at day 10. Dendrogram clustering and PCA separated MEFs and MEFs+MyoD into one group whereas iMPCs and MyoD+F/R/C treated MEFs into another group, with further subdivision in each respective group (Fig. 3A, Supplementary Fig. 4A). To corroborate this observation further, we identified the number of differentially expressed proteins (DEPs) between the various groups. In accordance with the hierarchical clustering, the percentage of DEPs was higher for MEFs subjected to MyoD+F/R/C treatment or iMPCs than MEFs subjected to MyoD condition alone, suggesting that MyoD+F/R/C manifests in more pronounced proteome changes (Fig. 3B). However, in comparison to parental MEFs we documented in all groups upregulation of several skeletal muscle associated proteins such as MYOD, DESM, DMD and TNNT1, albeit the proliferation markers PCNA and KI67 and myogenic proteins DEK and MUSK were significantly upregulated only in MyoD+F/R/C condition and stable iMPCs (Fig. 3C). We then analyzed the top 30 upregulated DEPs between each condition vs. MEFs and documented the upregulation of typical skeletal muscle proteins in all groups including MYH7, MYH8, TNNT2, ACTN3, and CASQ2 (Fig. 3D). Furthermore, a functional enrichment analysis of DEPs documented the upregulation of several skeletal muscle gene networks in all groups (Supplementary Fig. 4B). We then performed a direct comparison between MyoD+F/R/C and MyoD conditions at day 10 of reprogramming and documented 681 upregulated and 349 downregulated proteins out of 4097 total proteins (Fig. 3E). Of note, the most upregulated proteins in the MyoD+F/R/C vs. MyoD conditions were associated with skeletal muscle differentiation markers and signaling proteins such as CASQ1, MYOZ1, GDF8 (*Mstn*), IBP2, IBP5 and cell proliferation and chromatin regulators including KI67, CDN1C, DNMT1 and BRD7 (Fig. 3F). Following MyoD+F/R/C treatment we further observed significant enrichment of cell proliferation and DNA replication proteins (MCM2, MCM3, MCM4, MCM6, PCNA, KI67), chromatin regulators (DNMT1, UHRF1, MBD3, PARP1, SETD7), signaling proteins (IGF2, TGFB2, IBP2, IBP5, CREB1) and skeletal muscle-associated proteins (PAXI, PYGM, MYH4, DEK, BCAM, CASQ1, MYOZ1) (Fig. 3G).

**Fig. 3:**
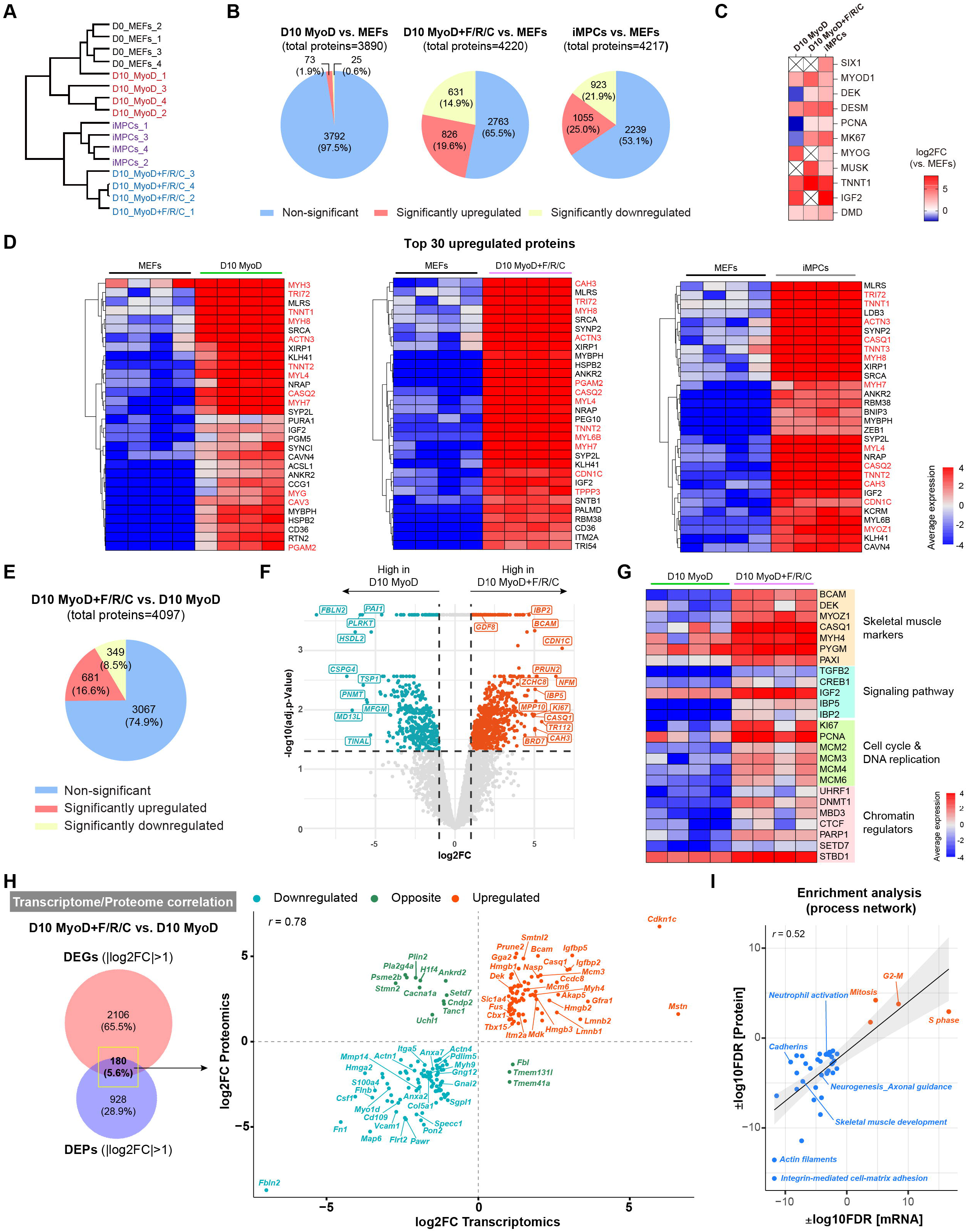
Proteome dynamics during iMPC formation. **(A)** Hierarchical clustering based on total proteome data. N = 4 cell lines per group. **(B)** Pie charts showing quantification of DEPs between the indicated cell lines. The significance threshold was set at |log2FC|>1 and adj. p-value<0.05. N = 4 cell lines per group. **(C)** A heatmap showing relative protein expression for canonical myogenic markers in the indicated cell lines. The calculated log2FC in each cell line vs. MEFs is shown. N = 4 cell lines per group. **(D)** Heatmaps based on proteome data displaying the top 30 upregulated proteins in the indicated cell lines vs. parental MEFs. The average expression is presented as gradient. N = 4 cell lines per group. Canonical myogenic differentiation-associated proteins are highlighted in red. **(E)** Pie chart showing quantification of DEPs between MEFs subjected to MyoD or MyoD+F/R/C conditions at day 10. The significance threshold was set as |log2FC|>1 and adj. p-value<0.05. N = 4 cell lines per group. **(F)** Volcano plot for DEPs between MEFs subjected to MyoD and MyoD+F/R/C conditions at day 10 of reprogramming. Significant DEPs (|log2FC|>1, adj.p-value<0.05) are shown as blue and red dots. N = 4 cell lines per group. **(G)** A heatmap of highly expressed proteins in MEFs subjected to MyoD+F/R/C conditions vs. MyoD condition at day 10 of reprogramming shown together with their respective cluster annotation. The average expression is presented as gradient. N = 4 cell lines per group. **(H)** Left-A Venn diagram showing the number of statistically significant DEGs and DEPs identified in MyoD+F/R/C vs. MyoD conditions at day 10 of reprogramming (|log2FC|>1, p-value<0.05). Right-scatterplot showing the correlation between the transcriptome and proteome of MyoD+F/R/C vs. MyoD conditions at day 10 of reprogramming. 180 overlapped DEGs / DEPs are projected on the plot, corresponding to the Venn diagram on the left. **(I)** Scatterplot showing the correlated Process networks that are enriched at the mRNA and protein levels in MyoD+F/R/C vs. MyoD treated MEFs at day 10. Enrichment analysis of DEGs / DEPs (|log2FC|>1, p-value<0.05) was performed using Metacore. Upregulated (in Red) and downregulated (in Blue) Process networks are shown using −log10(FDR) and log10(FDR), respectively.

Next, we performed an integrative analysis between the RNA-Seq and proteome datasets. We first inspected significant DEGs and DEPs in iMPCs compared to MEFs and recorded 443 shared genes/proteins with a high correlation (*r* = 0.81) (Supplementary Fig. 4C). Notably, enriched genes/proteins in iMPCs included skeletal muscle-related markers, chromatin regulators and signaling pathway factors, whereas MEFs were enriched for fibroblast-specific genes and proteins (Supplementary Fig. 4C). Given this observation, a similar integrative analysis was performed for MyoD+F/R/C vs. MyoD conditions at day 10 of reprogramming and a group of 180 shared DEGs and DEPs was identified (Fig. 3H). We noted a high correlation between the mRNA and protein levels (*r* = 0.78) and documented following F/R/C treatment a unique upregulation of skeletal muscle markers such as *Dek*, *Igfbp2*, *Igfbp5*, *Mstn*, *Bcam*, *Myh4* and *Casq1,* in addition to cell proliferation and DNA replication markers such as *Mcm3*, *Mcm6*, and *Nasp* (Fig. 3H). Lastly, we performed enrichment analysis for each group of upregulated and downregulated markers, thus revealing upregulation of “cell proliferation” associated networks and downregulation of “developmental processes” and “cell adhesion” associated networks following F/R/C treatment (Fig. 3I).

Collectively, the LC-MS analysis uncovered a unique expression of proteins which are significantly upregulated in iMPCs or MyoD+F/R/C condition. These proteins include muscle differentiation proteins, signaling factors, chromatin regulators and proliferation markers. The high expression of proteins such as MYOZ1, PYGM, MYH4 and DEK predominantly in the presence of F/R/C treatment suggests that iMPCs preferentially express mature muscle markers in comparison to myotubes produced via transdifferentiation by MyoD.

### Unique chromatin accessibility dynamics during formation of iMPCs

Chromatin accessibility in promoter regions is well-known to be associated with increased gene expression. Given the prominent differences at the mRNA and protein levels between MyoD and MyoD+F/R/C treatment, we reasoned that divergent chromatin accessibility dynamics in promoters of key myogenic genes may explain the observed differences in the transcriptome and proteome during transdifferentiation and reprogramming. To investigate this hypothesis, we performed an Assay for Transposase-Accessible Chromatin using sequencing (ATAC-Seq) of MEFs, MEFs subjected to MyoD or MyoD+F/R/C for 2 days, and an established iMPC clone. As expected from our previous multi-omics analyses, iMPCs were characterized by a distinct chromatin accessibility profile in comparison to MEFs, and surprisingly demonstrated a high percentage of annotated peaks around promoter regions (Fig 4A, B). Given this observation, we performed differential chromatin accessibility analysis in the gene promoter regions of canonical myogenic and fibroblast genes. In comparison to MEFs, we detected a decrease of chromatin accessibility in fibroblast-specific gene promoters such as *Thy1*, *Fbln2*, *Fbln5*, *Col1a1* and an increase in myogenic gene promoters including *Myh8*, *Dll1*, *Six1* and *Myod1* in all myogenic cell lines (Fig. 4C). In accordance with the transcriptome analysis, we also documented a preferential increase of chromatin accessibility in MyoD+F/R/C treated MEFs and iMPCs in multiple myogenic differentiation genes such as *Myf6*, *Mstn* and *Casq1* as well as progenitor genes such as *Sox8* and *Fgfr4* (Fig. 4C). Notably, open chromatin configuration in the promoters of muscle stem cell markers such as *Pax7*, *Myf5* and *Notch3* was solely documented in established iMPCs (Fig. 4C).

**Fig. 4:**
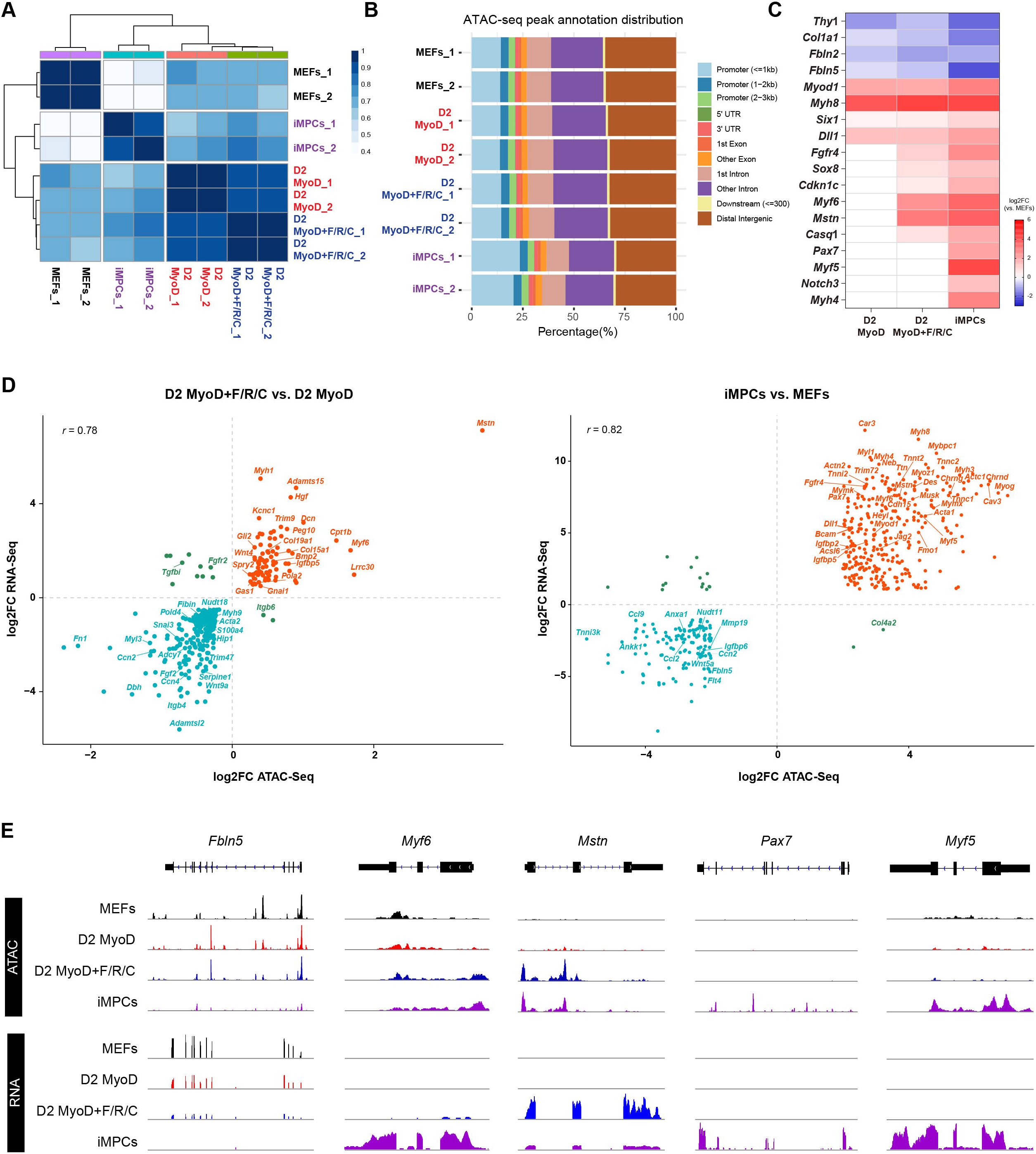
Chromatin accessibility dynamics unique to iMPC reprogramming. **(A)** Correlation matrix for the indicated samples based on ATAC-Seq for global chromatin accessibility. *Pearson’s correlation coefficient r* is displayed as a color gradient. **(B)** Distribution of annotated ATAC-Seq peaks across the respective genomic regions. **(C)** Heatmap showing differential chromatin accessibility of the indicated gene promoter regions (±1kb of TSS) compared to MEFs. Log2FC is shown via a color gradient and calculated vs. MEFs. Non-significant values (p-value > 0.01) are shown in white. N = 2 cell lines per group. **(D)** Scatterplot of selected genes showing the correlation between chromatin accessibility in promoter regions (±1kb of TSS) and bulk RNA-Seq in day 2 MyoD+F/R/C vs. MyoD treated MEFs (left) and iMPCs vs. MEFs (right). The genes with |log2FC| > 0.5 (left) and |log2FC| > 2 (right) are presented with p-value < 0.01. **(E)** IGV tracks of ATAC-Seq peaks and bulk RNA-Seq for the indicated genes.

To investigate further whether chromatin accessibility in promoter regions might be associated with differential gene expression, we performed an integrative analysis of ATAC-Seq and bulk RNA-Seq datasets between MyoD and MyoD+F/R/C treated MEFs at day 2, as well as established iMPCs and MEFs (Fig. 4D). For both comparisons, we noted a positive and high correlation between increased gene expression and an accessible and more “open” chromatin configuration (Fig. 4D). Of note, we detected an increase in both gene expression and chromatin accessibility in the promoter regions of myogenic genes that were preferentially expressed following F/R/C treatment at day 2 including *Mstn*, *Myf6*, *Myh1* and *Gas1* (Fig. 4D, E). Similarly, we recorded an increase of myogenic stem and differentiated cell markers unique to iMPCs in comparison to parental MEFs including *Pax7*, *Myf5*, *Bcam*, *Heyl*, *Mymk*, *Myh1*, *Myh4* (Fig. 4D, E). Collectively, our results demonstrate that as early as 2 days following MyoD or MyoD+F/R/C treatment of MEFs prominent chromatin accessibility changes can be detected in myogenic promoters, however F/R/C treatment further elicits preferential open chromatin in multiple key myogenic gene promoters, which may explain their unique transcriptome and proteome profiles at this time point.

### Cell types and differentiation trajectories in iMPCs uncovered by single cell RNA-Seq

The heterogeneity of iMPCs renders their entire molecular characterization challenging utilizing bulk multi-omics tools. To address this challenge and identify the underlying cell types that comprise the heterogeneous iMPC cultures, we performed single cell RNA-Sequencing (scRNA-Seq) of an iMPC clone. Utilizing unsupervised clustering^26^, we identified 9 distinct cell clusters with 3 divergent cell cycle states (Fig. 5A, B). Three of these clusters (1, 2, and 4) represented the Pax7^+^ stem cell fraction of iMPCs, and could be further separated into cycling (C1, 16.25%) or less cycling cell populations (C2, 14.88% or C4, 12.58%) (Fig. 5A-C). These clusters were also highly enriched for the myogenic stem and progenitor markers *Myf5, Dek, Mest, Plagl, Dbx1, Fzd4 and Sdc4* (Fig. 5C-D, Supplementary Fig. 5A). Notably, cells in cluster 5 (C5, 10.38%) expressed high levels of *Sox8*, *Myod1*, *Myog* and the canonical Notch ligand *Dll1,* thus representing committed progenitors, whereas late-stage differentiation markers including *Myf6*, *Mstn, Mymk*, *Mymx* and *Myh1* were present in cluster 6, thus representing fusion-competent myocytes (C6, 7.64%) (Fig. 5C, D, Supplementary Fig. 5A). Cells in cluster 7 (C7, 6.62%) may correspond to a more mature myofiber population given the high expression of *Myh1*, *Tnnt3*, and *Tnni1* but absence of *MyoD* or *Myog* (Fig. 5C, D). Surprisingly, three clusters (0, 3, and 8) highly expressed *Col1a1, Thy1, Pdgfra and Pdgfrb* markers which are indicative of connective tissue cells in the form of fibroblasts, mesenchymal cells or fibro-adipogenic progenitors (Fig. 5A, C). Lastly, we observed that pseudo gene expression of iMPCs from scRNA-Seq strongly correlated with gene expression of iMPCs from bulk RNA-Seq, and the connective tissue-like cell populations (C0, C3, C8) with bulk RNA-Seq of MEFs (Supplementary Fig. 5B, C). This correlation analysis collectively confirms that the cell identities revealed via scRNA-Seq faithfully represent the cell types we previously characterized using bulk RNA-Seq.

**Fig. 5:**
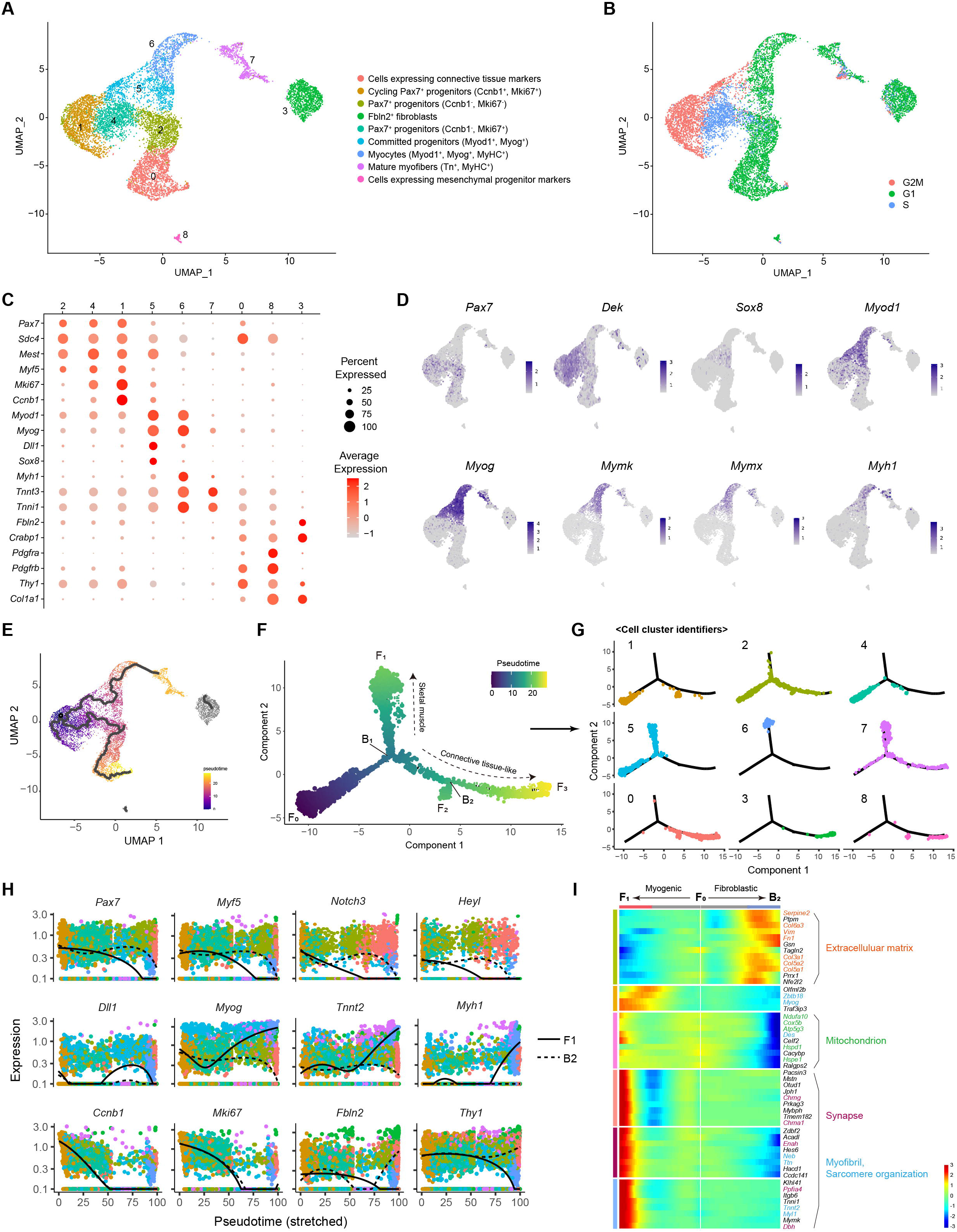
scRNA-Seq uncovers cell types and differentiation trajectories in iMPCs. **(A)** UMAP projection of the scRNA-Seq data showing 9,184 cells colored by clusters that form a stable iMPC clone. **(B)** UMAP projection colored by 3 main cell cycle states in an iMPC clone. **(C)** Dot plot showing the expression of the indicated genes in each cell cluster. The size of the dot indicates the percentage of cells expressing the indicated genes within the cluster. The color-coding scale represents the average gene expression level in all cells in a cluster. **(D)** UMAP projection showing all cells colored by the indicated myogenic markers based on scRNA-Seq data. **(E)** UMAP projection showing all cells colored by pseudotime as calculated from unsupervised single cell trajectory as reconstructed by Monocle 3. **(F-G)** Minimum spanning tree showing ordered cells based on semi-supervised single cell trajectory analysis reconstructed by Monocle2 and colored by either pseudotime (F) or cell cluster identifiers (G). The pseudotime trajectory initiates from the F_0_ root and bifurcates at the branch point B_1_ before proceeding towards two main cell fates (denoted as F_1_ and F_3_). **(H)** Plot showing the kinetics of the indicated genes in various cell types as a function of pseudotime (F) emanating from point F_0_ and proceeding towards F_1_ (Solid line, “myogenic cell fate”) or to B_2_ (Dashed line, “connective tissue cell fate”). Dots indicate cells colored by cell identifiers. **(I)** Heatmap based on scRNA-Seq data for the top 50 DEGs regulated at branch point B_1_ in (F). The individual gene expression level initiating at point F_0_ and proceeding towards a myogenic cells fate (F_0_->F_1_) is shown on the left and connective tissue cell fate (F_0_->B_2_) is shown on the right. Color-code gradient shows normalized gene expression level (Z-score) for each gene across all cells. The GO terms were annotated with DAVID v6.8 for each respective gene group shown using the same color coding.

To investigate whether iMPCs recapitulate a myogenic differentiation program *in vitro,* we performed a pseudotime lineage trajectory analysis using the Monocle package^27^. As first step, we performed an unsupervised trajectory analysis, which revealed two major cellular progressions, both emanating from the cycling Pax7^+^ progenitors (C1) (Fig. 5E). One branch further gave rise to several myogenic differentiated cell types (C5, C6, C7). Surprisingly, another branch emanated towards a less proliferative population of Pax7^+^ cells and gave rise to a connective tissue-like cell type (C0). Aside from these two main trajectories, C3 and C8 were not associated with an active lineage progression, suggesting that these may correspond to non-reprogrammed MEFs or other mesenchymal cells (Fig. 5E).

To validate the reproducibility of the trajectory analysis, we reconstructed a cellular progression in a semi-supervised manner. Using this approach, we determined two distinct differentiation branch points (denoted as B_1_ and B_2_) and three different cell fates (denoted as F_1_, F_2_, and F_3_) emanating from the root of trajectory F_0,_ which was mainly composed of Pax7^+^ progenitors (Fig. 5F, G Supplementary Fig. 5D). The B_1_ point represents a clear bifurcation into two distinct reprogramming routes, F_1_ and F_3_ (Fig. 5F). Notably, the progression from B_1_ to F_1_ mainly involves cells that express committed and differentiation myogenic genes (C5, C6, C7), whereas the progression from B_1_ to F_3_ consists of connective tissue-like cells (C0, C3, C8) (Fig. 5F, G). Lastly, we investigated the individual gene level expression in the two differentiation routes emanating from B_1_. First, stem and progenitor cell markers such as *Pax7* and *Myf5* were highly expressed at the start of the pseudotime and were rapidly downregulated as they progressed to F_1_ (Fig. 5H, Supplementary Fig. 5E). Additionally, the committed progenitor genes *Myod1* and *Myog* were only upregulated from B_1_ to F_1_, in conjunction with appearance of differentiation markers such as *Myh1*, *Tnni1*, *Tnnt2*, *Tnnt3*. Remarkably, *Dll1* and *Sox8* expressing cells were detected in the middle of the branch leading to F_1_, further suggesting that this cell population represent committed myoblasts progressing to become differentiated skeletal muscle cells. In addition, the cell cycle proliferation markers *Ccnb1* and *Mki67* were significantly downregulated in both routes, whereas the connective tissue markers *Fbln2*, *Thy1*, *Col1a1*, *Pdgfra* and *Pdgfrb* were only upregulated in the path leading to B_2_. Lastly, differential gene expression analysis between the two routes confirmed our observation that highly expressed genes in B_2_ were annotated with *Extracellular matrix,* whereas the genes in F_1_ were annotated with *Synapse*, *Myofibril and sarcomere organization,* thus reinforcing the notion that cells in B_2_ are skeletal muscle cells (Fig. 5I).

In summary, utilizing scRNA-Seq we identified the various cell populations that comprise the heterogeneous iMPC cultures, demonstrating the presence of cycling *Pax7*^+^ stem cells, *Myod1*^+^/*Myog*^+^ committed progenitors and *Myf6*^+^/*Myh1*^+^ differentiated cells, in addition to connective tissue-like cells. Moreover, we reconstructed the myogenic program in an iMPC clone using a pseudotime trajectory analysis and delineated the differentiation route from activated satellite-like cells into committed progenitor cells and differentiated muscle cells. This analysis also underscored an alternative differentiation route into a connective tissue-like cell type, cautiously suggesting that iMPCs may harbor a bipotential differentiation propensity.

### The molecular landscape of purified Pax7-nGFP^+^ iMPCs in comparison to myoblasts

The scRNA-Seq analysis uncovered a unique cell population consisting of Pax7^+^ stem cells, however utilizing bulk RNA-Seq we could only detect a fraction of the myogenic stem cell-associated genes in this population due to the heterogeneity of iMPC cultures. Furthermore, utilizing mass spectrometry we did not detect significant levels of satellite cell and myoblast-associated proteins in bulk iMPCs. This could be due to technical limitations in utilizing LS-MS to detect lowly expressed proteins in the form of stem cell-specific transcription factors due to the high expression of structural and signaling proteins emanating from the multinucleated myofibers of iMPCs.

To address this limitation and characterize the Pax7^+^ stem cell population in depth, we opted to FACS-purify Pax7-nGFP^+^ cells from iMPC clones and molecularly compare them to FACS-purified Pax7-nGFP^+^ myoblasts by bulk RNA-Seq and LC-MS (Fig. 6A). The expression levels of canonical satellite cell markers such as *Pax7* and *Six4* were higher in Pax7-nGFP^+^ iMPCs and Pax7-nGFP^+^ myoblasts, whereas the expression of skeletal muscle differentiation genes such as *Myh1*, *Myh4*, *Myh8* and *Casq1* was higher in bulk iMPCs, indicating that the stem cell sorting strategy from bulk iMPCs was successful (Fig. 6B). Next, to identify pathways unique to the stem cell subsets of iMPCs, we performed a pathway enrichment analysis between bulk iMPCs and Pax7-nGFP^+^ iMPCs, thus revealing gene categories that were highly enriched in bulk iMPCs and associated with differentiated muscle cells (Fig. 6C, D). In contrast, “cell proliferation” and “metabolism/chromatin modulation” associated pathways were highly enriched in Pax7-nGFP^+^ iMPCs in addition to the *Notch* and *Hedgehog* signaling pathways (Fig. 6C, D).

**Fig. 6:**
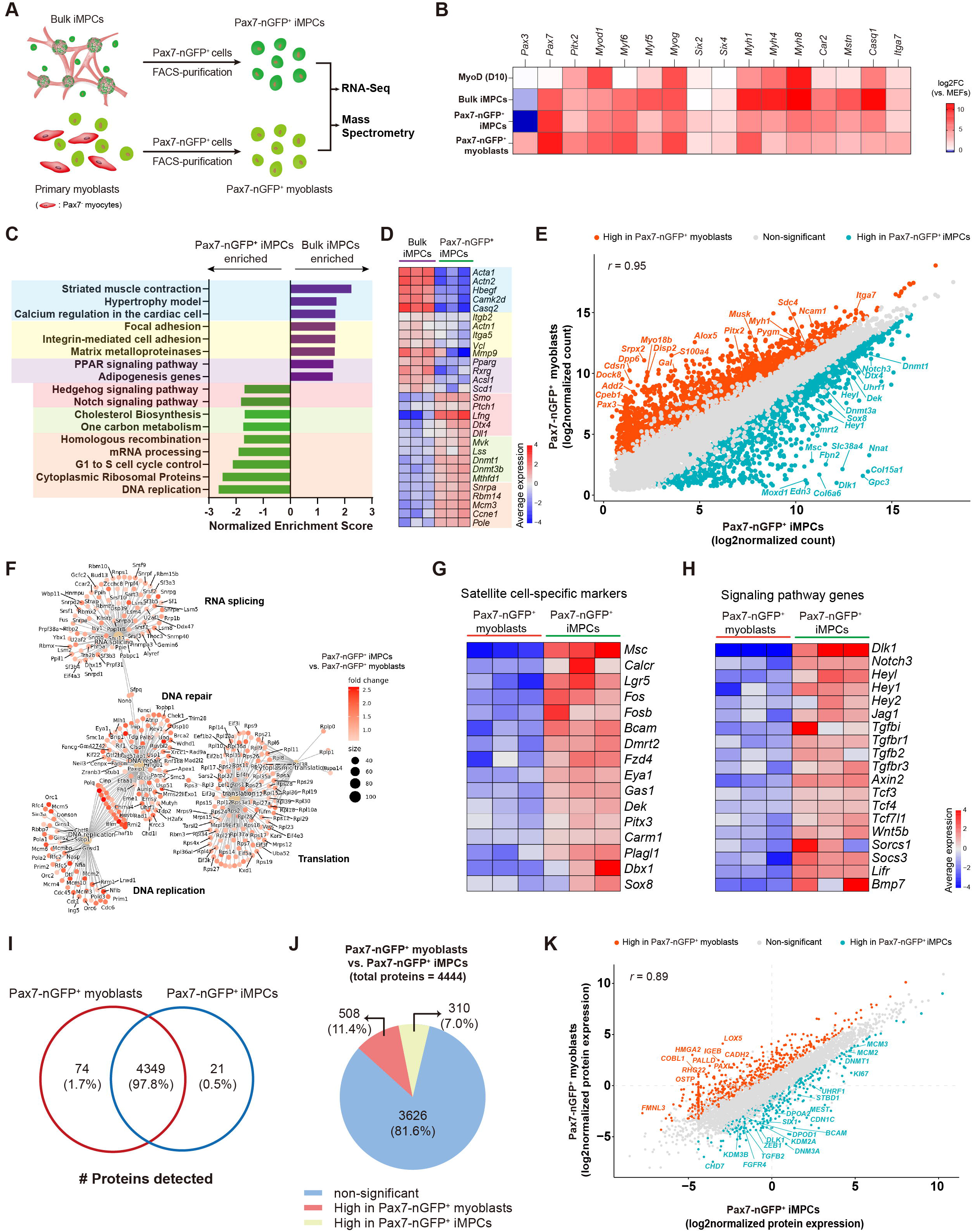
Molecular characterization of FACS-purified Pax7-nGFP^+^ iMPCs and myoblasts. **(A)** schematic illustrating the strategy to FACS-purify Pax7-nGFP^+^ cells from heterogeneous iMPCs or primary myoblasts prior to bulk RNA-Seq and proteomic analyses. Heatmap based on bulk RNA-Seq data showing the expression level of skeletal muscle related genes in the indicated samples. Log2FC of respective cell types vs. MEFs is shown. N = 3 cell lines per group. **(C)** Pathway enrichment analysis using GSEA between bulk iMPCs and Pax7-nGFP^+^ iMPCs. **(D)** Heatmap based on bulk RNA-Seq dataset demonstrating the expression level of the indicated genes between Pax7-nGFP^+^ iMPCs and bulk iMPCs. Each gene group is associated with the respective pathway annotation in (C). The average expression is presented as gradient. N = 3 cell lines per group. **(E)** Scatter plot showing gene expression based on bulk RNA-Seq data between Pax7-nGFP^+^ iMPCs and Pax7-nGFP^+^ myoblasts. Log2 of normalized read counts is presented. Red and blue dots denote upregulated genes (log2FC>1) in Pax7-nGFP^+^ myoblasts or Pax7-nGFP^+^ iMPCs, respectively. Gray dots indicate non-significant genes between the two cell types. N = 3 cell lines per group. **(F)** Over Representation Analysis (ORA) of the gene sets that were significantly enriched (log2FC > 0.5, p-value < 0.01) in Pax7-nGFP^+^ iMPCs in comparison to Pax7-nGFP^+^ myoblasts. The size indicates the number of genes involved in the annotation and the color-coding scales the fold change. **(G-H)** Heatmaps based on bulk RNA-Seq data of candidate genes associated with satellite cell markers (G) and signaling pathways (H) that are highly expressed in Pax7-nGFP^+^ iMPCs. The average expression is presented via a gradient. N = 3 cell lines per each group. **(I)** A Venn diagram based on proteome data showing the total number of proteins detected in Pax7-nGFP^+^ myoblasts and Pax7-nGFP^+^ iMPCs. N = 4 cell lines per group. **(J)** Number of DEPs in Pax7-nGFP^+^ myoblasts vs. Pax7-nGFP^+^ iMPCs. The significance threshold was set as |log2FC|>1 and adj. p-value<0.1. N = 4 cell lines per group. **(K)** Scatter plot based on log2 normalized protein expression between Pax7-nGFP^+^ iMPCs and Pax7-nGFP^+^ myoblasts. N = 4 cell lines per group.

The FACS-purification strategy further allowed us to directly compare Pax7-nGFP^+^ iMPCs to Pax7-nGFP^+^ myoblasts. As expected, the two cell types were transcriptionally similar (*r* = 0.95) (Fig. 6E). However, we also documented statistically significant DEGs which included unique satellite cell markers in addition to proliferation and chromatin regulators in Pax7-nGFP^+^ iMPCs (Fig. 6E). Further, an Over Representation Analysis (ORA) showed divergent transcriptional categories unique to Pax7-nGFP^+^ iMPCs or myoblasts (Fig. 6F, Supplementary Fig. 6A). Strikingly, we detected high expression of activated satellite cell markers in Pax7-nGFP^+^ iMPCs that were not detectable or lowly expressed in Pax7-nGFP^+^ myoblasts (Fig. 6G). These included *Calcr*, *Musculin* (*Msc*), *Lgr5*, *Fos*, *Dmrt2*, *Fzd4*, *Gas1*, *Dek*, *Pitx3*, *Carm1*, *Sox8*, *Dbx1* and *Plagl*, many of which have been recently reported to regulate *in vivo* muscle regeneration and satellite cell activation^28–36^(Fig. 6G). Moreover, genes associated with critical pathways for satellite cell activation and proliferation, including Notch, TGF-β, JAK-STAT and WNT were significantly upregulated in Pax7-nGFP^+^ iMPCs (Fig. 6H). Lastly, we documented in Pax7-nGFP^+^ iMPCs significant enrichment for chromatin remodelers, including *Tet1*, *Tet3*, *Dnmt1*, *Dnmt3a*, *Dnmt3b* and *Uhrf1,* members of the transcription factor families Myc, KLF and BEX and a plethora of cell proliferation and DNA replication markers in comparison to myoblasts (Supplementary Fig. 6B, C, D).

To examine whether the transcriptional differences could be further detected at the protein level, we opted to perform LC-MS on Pax7-nGFP^+^ iMPCs and Pax7-nGFP^+^ myoblasts. Using this approach, we detected 4444 proteins, of which 4349 were detected in both cell types (Fig. 6I). From this protein group, 508 and 310 proteins were differentially expressed in Pax7-nGFP^+^ myoblasts and Pax7-nGFP^+^ iMPCs, respectively (Fig. 6J). A comparison of DEPs between the two cell types documented transcription factors and chromatin regulators which were significantly more expressed in Pax7-nGFP^+^ iMPCs in comparison to Pax7-nGFP^+^ myoblasts including BCAM, CHD7, SIX1, DNM3A, ZEB1, FGFB2, FGFR4, KDM2A and the cell proliferation markers KI67, MCM2 and MCM3 (Fig. 6K). In summary, using a transgenic Pax7 reporter we successfully purified and characterized the transcriptome and a portion of the proteome of Pax7-nGFP^+^ iMPCs and myoblasts. This analysis established a unique expression signature in Pax7-nGFP^+^ iMPCs which is reminiscent of activated satellite cells and distinct from primary myoblasts.

### The Notch pathway is critical for iMPC formation and self-renewal

Our transcriptional analysis thus far underpinned the involvement of the Notch pathway during iMPC formation. As such, we hypothesized that the Notch pathway is critical for iMPC formation as previously reported for proliferation of endogenous satellite cells *in vivo*^37–39^. Indeed, we detected early and rapid upregulation of *Notch1*, *Notch3*, *Hey1* and *Heyl* only in MyoD+F/R/C during the reprogramming course (Fig. 7A). To test whether the Notch pathway is critical for iMPC formation, we treated MEFs undergoing MyoD or MyoD+F/R/C conversion with DAPT, an inhibitor of Notch target γ-secretase^40^. We documented complete lack of iMPC formation and cell proliferation at day 10 of MyoD+F/R/C+DAPT treatment in comparison to MyoD+F/R/C treated cells, albeit multinucleated myotubes formed in both conditions (Fig. 7B).

**Fig. 7:**
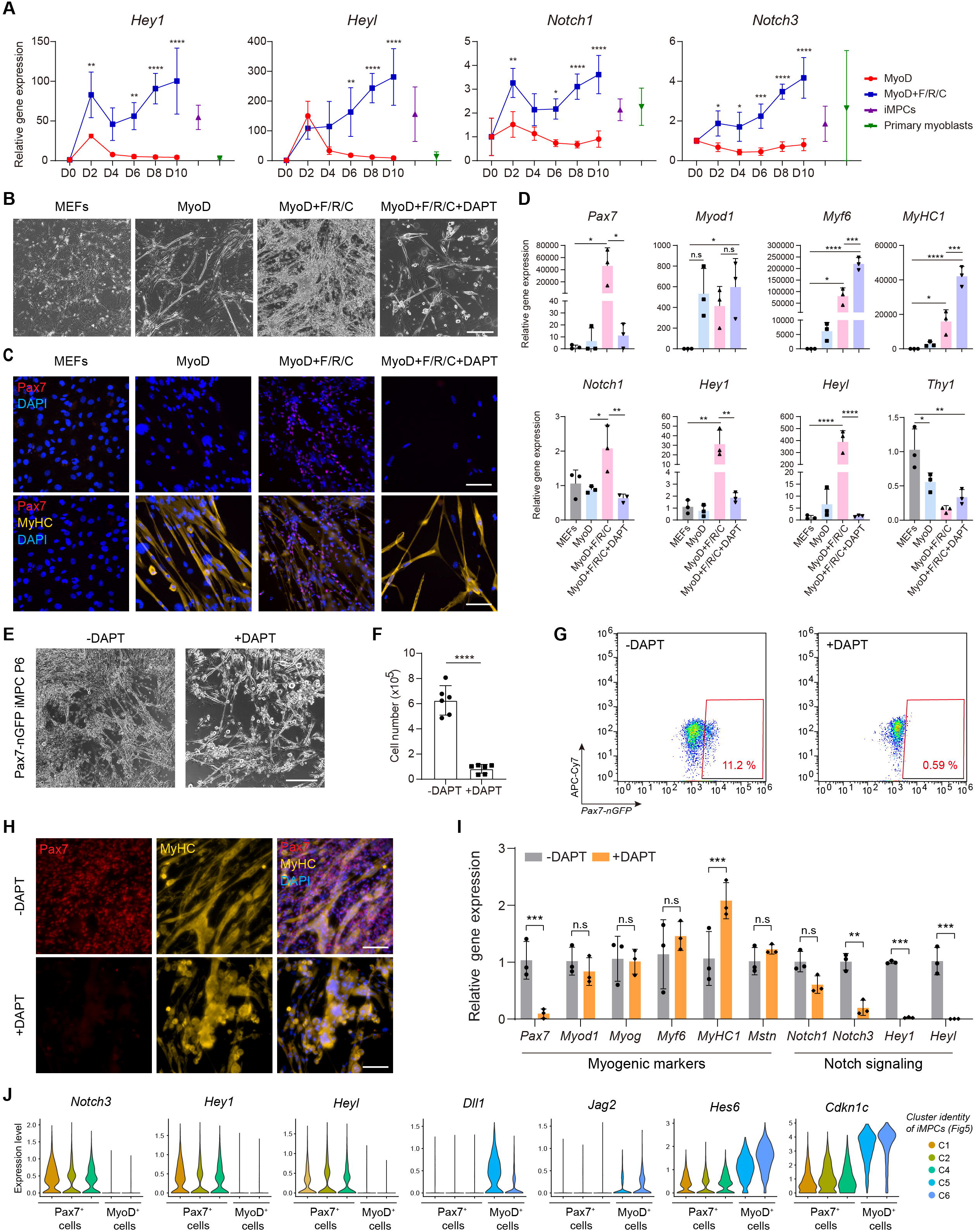
Notch signaling is indispensable for the formation and maintenance of iMPCs. **(A)** Expression dynamics based on bulk RNA-Seq data for the indicated Notch pathway associated genes during MyoD or MyoD+F/R/C reprogramming course. Established iMPCs and primary myoblasts served as positive controls. Relative gene expression was calculated by normalizing the RPKM values of each sample to that of parental MEFs (D0 MEFs). N = 3 cell lines per group. Statistical significance was determined by two-way ANOVA between the conditions at each time point (*p<0.05, **p<0.01, ***p<0.001, ****p<0.0001). **(B)** Representative bright-field images of MEFs subjected to MyoD, MyoD+F/R/C and MyoD+F/R/C+DAPT treatment at day 10 of reprogramming. Scale bar, 400 μm. **(C)** Representative immunofluorescence images of MEFs subjected to the listed conditions at day 10 of reprogramming and stained for Pax7, MyHC and DAPI. Scale bar, 100 μm. **(D)** qRT-PCR analysis for the indicated genes and conditions. Relative gene expression is shown as means□±□S.D. N = 3 cell lines per each group. Statistical significance was determined by one-way ANOVA (*p<0.05, **p<0.01, ***p<0.001, ****p<0.0001, n.s=non-significant). **(E)** Representative bright-field images of iMPCs (P6) treated or non-treated with DAPT for 5 consecutive days. Scale bar, 400 μm. **(F)** Quantification of the total number of cells in iMPCs treated or non-treated with DAPT for 5 consecutive days. Statistical significance was determined by a two-tailed unpaired *t*-test (****p<0.0001). N= 2 experimental repeats of 3 cell lines per group. **(G)** Flow cytometry analysis for Pax7-nGFP iMPCs treated or non-treated with DAPT for 5 days. **(H)** Representative immunofluorescence images for Pax7, MyHC and DAPI in iMPCs treated with and without DAPT for 5 consecutive days. Scale bar, 100 μm. **(I)** qRT-PCR for selective genes in DAPT-treated iMPCs for 5 consecutive days. The data is shown as meansL±LS.D. N = 3 cell lines per group. Statistical significance was determined by one-way ANOVA (**p<0.01, ***p<0.001, n.s = non-significant). **(J)** Violin plots showing the average expression of the indicated genes in the respective clusters based on scRNA-Seq data from Fig 5. Note presence of canonical Notch-related markers in Pax7^+^ iMPCs (clusters 1, 2, 4) as well as typical Notch ligands and inhibitors in their differentiated progeny (clusters 5, 6).

To unequivocally confirm the lack of iMPC formation following DAPT treatment, we assessed the expression of Pax7 and MyHC in MEFs subjected to MyoD+F/R/C+DAPT treatment for 10 days. Unlike MyoD+F/R/C condition, we did not detect Pax7^+^ cells in MyoD+F/R/C+DAPT condition, albeit MyHC^+^ myotubes did form (Fig. 7C). In accordance with the RNA-seq data, qRT-PCR revealed that the expression of *Pax7*, *Notch1*, *Hey1* and *Heyl* was higher in the MyoD+F/R/C than MyoD condition, however DAPT treatment abrogated the expression of these genes to levels reminiscent of MEFs or MEFs+MyoD (Fig. 7D). DAPT treatment did not alter the expression of other myogenic markers such as *Myf5*, *Myod1, Mstn* and *Myog* as well as the fibroblast-specific marker *Thy1*, suggesting that Notch inhibition preferentially affects the Pax7^+^ stem cell subset (Fig. 7D, Supplementary Fig. 7).

We next assessed the effect of Notch inhibition on the self-renewal of stable iMPC clones. To this end, we cultured iMPCs in the presence of DAPT for 5 consecutive days, observing a marked decrease in cell density and complete depletion of Pax7-nGFP^+^ cells (Fig. 7E-G). We also confirmed the absence of Pax7^+^ cells via immunofluorescence and noted that MyHC^+^ myofibers were detected in both conditions (Fig. 7H). To investigate further gene regulation following DAPT treatment, we conducted qRT-PCR for myogenic and Notch-related genes in DAPT treated and non-treated iMPCs. This analysis revealed a marked reduction of Pax7 expression in concert with downregulation of *Hey1*, *Heyl*, *Notch1* and *Notch3* in DAPT treated iMPCs, whereas *Myod1*, *Myog*, and *Mstn* were expressed in similar fashion (Fig. 7I).

Next, we analyzed the expression level of Notch pathway-associated genes in the various cell subsets comprising a stable iMPC clone utilizing the scRNA-Seq dataset (Fig 5). In accordance with the bulk RNA-Seq analysis of FACS-purified Pax7-nGFP^+^ iMPCs, we documented Pax7^+^ cells which highly expressed the Notch receptors *Notch3* and *Notch1*, in addition to their canonical downstream targets *Heyl* and *Hey1* (Fig. 7J). Strikingly, the cell subset of Sox8^+^/MyoD^+^ progenitor cells and MyHC^+^/Myog^+^ differentiated cells expressed the canonical Notch ligands *Dll1*, *Jag2* and *Dlk1* as well as the Notch inhibitors *Cdkn1c* and *Hes6* (Fig. 7J). This observation suggests a crosstalk via the Notch pathway between the stem cells and progenitor / differentiated cell subsets in iMPCs.

Collectively, we surmise that activation of the Notch pathway is a unique feature of Pax7^+^ cell formation during MyoD+F/R/C reprogramming. DAPT-mediated Notch pathway inhibition during MyoD+F/R/C reprogramming and in iMPC clones precludes the formation of Pax7^+^ cells and derails their self-renewal. However, myocytes and myotubes can still form following DAPT treatment, suggesting that Notch inhibition specifically prevents the formation of myogenic stem cell during iMPC reprogramming, and their maintenance in established iMPC clones. Lastly, stable iMPCs contain both Pax7^+^ stem cells that express canonical Notch markers in addition to their differentiated progeny which express Notch-associated ligands and inhibitors, thus recapitulating *in vitro* their expression during *in vivo* muscle regeneration.

## Discussion

Direct lineage reprogramming of somatic cells into multipotent stem or progenitor cells affords an attractive approach to generate desired cell types for basic research or therapeutic applications. This approach entails several advantages in comparison to canonical transdifferentiation, which typically involves direct conversion of one differentiated cell type into another. Namely, directly reprogrammed progenitors may exhibit self-renewal and multipotency, rendering them more attractive for cell-based therapies. However, most studies to date have reported on protocols to directly transdifferentiate cells, and only a handful of studies documented direct conversion of somatic cells into multipotent progenitors^9^. Furthermore, numerous studies have characterized the molecular transitions and mechanisms governing transdifferentiation, yet very few interrogated how the molecular landscape metamorphoses during direct conversion into multipotent progenitor cells^9^.

In this study we set out to address this objective using the skeletal muscle lineage as a model system. To this end, we dissected the molecular changes that accompany fibroblast conversion into myogenic stem and progenitor cells by way of sustained MyoD overexpression in concert with administration of the three small molecules Forskolin, RepSox and CHIR99210 (F/R/C)^20^. Utilizing multi-omics approaches, we contrasted this lineage conversion to that of canonical MyoD-mediated transdifferentiation and delineated an array of genes, proteins and signaling pathways that are unique to each cell fate conversion. We demonstrated that reprogramming to iMPCs occurs via a gradual, stepwise reprogramming process, whereas transdifferentiation into myotubes is typically fast and direct. Additionally, we report that the two cell conversions share phenotypical characteristics, including rapid upregulation of several skeletal muscle differentiation genes, albeit are also distinct, as only F/R/C administration manifests a robust upregulation of satellite cell genes and key skeletal muscle differentiation markers via rewiring of the chromatin in promoters of key myogenic genes. We then further compared the transcriptomes and proteomes of FACS-purified Pax7-nGFP^+^ iMPCs to that of Pax7-nGFP^+^ primary myoblasts and documented their congruent and divergent molecular traits. Lastly, we identified Notch as a molecular pathway that is absolutely essential for the formation of iMPCs in addition to governing their self-renewal.

Several observations emanate from this study. One notable finding pertains to the molecular comparison of Pax7-nGFP^+^ iMPCs to that of Pax7-nGFP^+^ myoblasts. These two cell types share molecular attributes at the mRNA and protein levels, however Pax7-nGFP^+^ iMPCs also uniquely expressed a cohort of genes which are indicative of satellite cell activation and proliferation *in vivo,* in addition to increased expression of cell cycle regulators and unique signaling pathways. We postulate several reasons which may account for these disparities. First, the Pax7-nGFP^+^ iMPCs are cultivated in the vicinity of neighboring committed progenitors, connective tissue-like cells and a highly contractile myofiber network, in contrast to primary myoblasts which are typically cultured as dispersed mononucleated cells that are passaged before fusion into myotubes. As such, the heterogeneity of iMPCs may recapitulate the microenvironment activated satellite cells encounter during skeletal muscle regeneration *in vivo*. Moreover, recent works have reported a crosstalk between resident muscle cells, myofibers and satellite cells during homeostasis and regeneration^41–44^. Similarly, signaling molecules secreted from the multinucleated myofibers of iMPCs may affect the gene expression of Pax7-nGFP^+^ iMPCs, potentially rendering them more akin to activated satellite cells. An additional explanation for the discrepancies between iMPCs and myoblasts involves the different culture conditions used to cultivate these cells. Whereas primary myoblasts were cultured on matrigel coated plates and in medium containing bFGF and high serum^45^, the iMPCs were cultured directly on plastic dishes using cell medium containing Serum Replacement, Serum, bFGF and most notably F/R/C. These cell media supplements may affect molecular attributes and elicit repression or activation of multiple myogenic genes and signaling pathways. In accordance with this hypothesis, exposure of dissociated skeletal muscle tissue fragments to iMPC media containing F/R/C treatment facilitated the formation of heterogeneous cultures consisting of myogenic progenitor cells and a contractile myofiber network that resemble fibroblast-derived iMPCs^20^.

Direct reprogramming using MyoD+F/R/C triggers extensive molecular transformations. One notable pathway that was uniquely upregulated following F/R/C treatment is Notch. This pathway has been extensively studied in the context of satellite cell quiescence and activation, and was shown to be critical for their self-renewal^32,46–48^. In this study, we demonstrate that this pathway is redundant for myotube formation, however, is critical for the generation of Pax7^+^ iMPCs and their maintenance. Moreover, we documented a high expression of Notch pathway-associated genes such as *Notch3*, *Heyl* and *Hey1* in the Pax7^+^ subset of iMPCs, whereas Notch ligands such as *Dll1*, *Jag2* and *Dlk1* were expressed in downstream differentiated myoblasts and myocytes. This observation is reminiscent of the Notch receptor / ligand interaction *in vivo*, suggesting a potential role for the differentiated cell subset of iMPCs in triggering elevated Notch levels in the Pax7^+^ cell subset. It will be of further interest to compare the levels of the Notch intra-cellular domain (NICD) in iMPCs in addition to culturing them on Notch ligands to assess whether such treatment may elicit a higher expression of the Notch pathway genes and increase the self-renewal of Pax7^+^ cells as previously shown for satellite cells^49,50^. How each small molecule exerts its effect during reprogramming or supports the proliferation of established iMPC clones is yet to be fully explored. Previous works have established the effect of Forskolin on enhancing myoblast proliferation and engraftment capacities in mice, and TGF-β inhibition was recently shown to promote myoblast fusion^51–53^. When administered together, these molecules have also been reported to enhance satellite cell quiescence in a 3D skeletal muscle tissue bioconstruct^54^. It is of interest to further explore their individual effects during reprogramming and whether F/R/C administration may augment the propensity of satellite cells to repair skeletal muscles *in vivo*.

To date, the conversion of somatic cells into iMPCs has only been reported from WT fibroblasts. As iMPCs recapitulate a unique myogenic differentiation program *in vitro*, it will be of interest to attempt their establishment from fibroblasts of murine models for muscular dystrophies. Namely, generation of iMPCs from a mouse model for Duchenne muscular dystrophy can provide a population of proliferative progenitors and contractile myofibers that lack dystrophin expression, affording a novel mean to model disease pathology *in vitro*, a platform for drug screens or a cell source for autologous cell therapy following genetic correction. Finally, several recent studies reported on direct conversion of mouse somatic cells into iMPCs using a variety of transcription factors and small molecules^18–20^. Reports on the conversion of human fibroblasts directly into myogenic stem and progenitor cells with satellite cell attributes and robust engraftment capacities are still lacking, and further work is highly warranted to establish such cell lines. We envision that with success, human iMPCs could serve as a complimentary toolbox in the skeletal muscle field for basic research, disease modeling and as a potential source for cell-based therapies.

## Acknowledgments

We wish to thank all members of the Regenerative and Movement Biology laboratory for their constructive comments and feedback. We are grateful to Dr. Shahragim Tajbakhsh for providing the Pax7-nGFP mouse strain and Dr. Katrien De Bock for fruitful discussions. We acknowledge the use of the Functional Genomics Center Zurich (FGCZ) facility and infrastructure and are grateful to Lennart Opitz and Laura Kunz for their assistance with data analysis of RNA sequencing and LC-MS.

## Funding

Eccellenza Grant of the Swiss National Science Foundation PCEGP3_187009 (OBN)

The Good Food Institute Foundation of the Swiss National Science Foundation 1-007082 (OBN)

The Novartis Foundation for Medical-Biological Research of the Swiss National Science Foundation 1-006449 (OBN)

The Helmut Horten Foundation of the Swiss National Science Foundation 1-007211 (OBN) The National Centre of Competence in Research (NCCR) Robotics of the Swiss National Science Foundation 5-29626 (OBN)

The European Research Council (ERC) under the European Union’s Horizon 2020 research and innovation program 803491 (FVM)

## Author contributions

Experiment design: IK, AG, OBN

Conceptualization: IK, OBN

Investigation: IK, AG

Methodology: IK, NB, LH

Supervision: OBN, FVM

Writing – original draft: OBN, IK

Writing – review & editing: IK, AG, NB, LH, FVM, OBN

## Competing interests

The authors declare that they have no competing interests.

## Data and materials availability

Plasmids generated in this study can be obtained from the Lead Contact or VectorBuilder (ID VB170530-1031pbc, ID VB181022-1110vfj). The bulk RNA-Seq and scRNA-Seq datasets generated in this study are available in Gene Expression Omnibus (GEO) repository (GSE169053 and GSE169054, respectively).

## Methods

### Animals

Mice carrying a Pax7-nGFP reporter (*Tg:Pax7-nGFP/C57BL6;DBA2*) were previously generated^55^. All mice used in this study were housed with 3-4 littermates and maintained under specific-pathogen-free (SPF)-like conditions. Mice were fed standard food and water and handled in accordance with the Swiss Federal Law on Animal Protection. All animal procedures were approved by the Zürich Cantonal Animal Welfare Committee (license ZH108/18).

### Plasmid construction

The plasmids used in this study were generated by VectorBuilder. *LV-EF1*α*-rtTA3/PGK-Neomycin* and *LV-tetO-MyoD/PGK-Puromycin* denote the respective plasmids: *pLV[Exp]-Neo-EF1A>Tet3G* (VectorBuilder, VB170530-1031pbc) and *pLV[Tet]-Puro-TRE3G>mMyod1[NM_010866. 2]* (VectorBuilder, VB181022-1110vfj).

### Virus production and storage

About 60-70% confluent HEK-293T cells were prepared in a 15 cm culture plate. For plasmid transfection, Δ8.9 (16.5 μg), VSV-G envelope (11 μg) and 22 μg of the target plasmid (either *pLV[Exp]-Neo-EF1A>Tet3G* or *pLV[Tet]-Puro-TRE3G>mMyod1*) were mixed in 150 mM NaCl at a final volume of 1 mL, followed by 10 mins incubation with 1 mL of 2 mg/mL Polyethylenimine (PEI) (Polysciences, POL23966-1). As next step, HEK-293T cells were incubated with a DNA/PEI mixture solution. After 1 day, the DNA/PEI mixture was replaced with fresh medium. At days 2 and 3, medium containing virus was collected and filtered through a 0.45 μm syringe filter (Corning, 431220). For virus precipitation, filtered medium was incubated with 5X PEG-it solution (System Biosciences, LV825A-1) overnight and centrifugated at 1500 xg for 30 mins at 4°C. The virus pellet was then re-suspended in PBS (Thermo Fisher Scientific, 10010015) containing 25 mM HEPES buffer (Thermo Fisher Scientific, 15630056) in 1/10 to 1/100 of original volume and stored at −80°C.

### Generation of Thy1^+^ reprogrammable MEFs (Rep-MEFs)

Mouse embryonic fibroblasts (MEFs) were isolated from *Pax7-nGFP* mice and cultured in ‘MEF medium’ containing high glucose DMEM (Thermo Fisher Scientific, 41966029) supplemented with 10% FBS (Thermo Fisher Scientific, 10270106), 1% GlutaMAX (Thermo Fisher Scientific, 35050061), 1% non-essential amino acids (Thermo Fisher Scientific, 11140050), 1% Pen-Strep (Thermo Fisher Scientific, 15140122) and 0.1% 2-Mercaptoethanol (Thermo Fisher Scientific, 21985023). To generate dox-inducible reprogrammable MEFs (Rep-MEFs), cells were passaged in a 6 well plates and once confluent transduced with *LV-EF1*α*-rtTA3/PGK-Neomycin* plus *LV-tetO-MyoD/PGK-Puromycin* at 1:1 ratio and supplemented with 6 μg/mL of polybrene (Sigma-Aldrich, TR-1003-G). Transduced MEFs were expanded and selected by sequential antibiotics treatment with MEF medium containing either 1 mg/mL of G-418 solution (Sigma-Aldrich, 4727878001) or 1 μg/mL of Puromycin (Thermo Fisher Scientific, A1113803) for a total of 4 days. To establish a homogenous population of fibroblasts from Rep-MEF cultures, we FACS-purified these cultures with an antibody recognizing the fibroblast-specific surface marker CD90.2 (Thy 1.2) (Thermo Fisher Scientific, 48-0902-80). Cells were FACS-purified or analyzed using an SH800S FACS-Sorter (Sony Biotechnologies).

### Reprogramming of Thy1^+^ Rep-MEFs

Approximately 2.5×10^5^ – 3.0×10^5^ Thy1^+^ MEFs were seeded onto 6 well plates and treated by means of the following conditions 1 day post seeding. For MyoD condition, Thy1^+^ Rep-MEFs were cultured with 2 μg/mL of doxycycline (dox) (Sigma-Aldrich, D9891) in ‘iMPC medium’ containing KnockOut DMEM (Thermo Fisher Scientific, 10829018) supplemented with 10% KnockOut Serum Replacement (Thermo Fisher Scientific, 10828028), 10% FBS, 1% GlutaMAX, 1% non-essential amino acids, 1% Pen-Strep, 0.1% 2-Mercaptoethanol and 10 ng/mL basic FGF (R&D Systems, 233-FB). For MyoD+F/R/C condition, Thy1^+^ Rep-MEFs were cultured in ‘iMPC medium’ containing 2 μg/mL of dox and administrated three small molecules: 5 μM of Forskolin (F) (R&D Systems, 1099/50), 5 μM of RepSox (R) (R&D Systems, 3742/50) and 3 μM of CHIR99021 (C) (R&D Systems, 4423/50).

### Establishment and maintenance of stable iMPC clones

To establish iMPC clones, Thy1^+^ Rep-MEFs were cultured in MyoD+F/R/C condition for 10 days, followed by another 2-3 days of culture in iMPC medium that contains only Forskolin, RepSox and CHIR99210 (without dox). For maintenance of stable clones, P0 iMPC clones were trypsinized and further expanded in iMPC medium containing the three small molecules.

### Satellite cell isolation

Whole body skeletal muscles were harvested from *Pax7-nGFP* mice and minced thoroughly with surgical scissors. PBS was added to minced muscles, followed by centrifugation at 350 xg for 2 mins. Cell pellets were incubated with 0.2% Collagenase type 2 (Thermo Fisher Scientific, 17101015) in high glucose DMEM for 90 mins in a 37°C shaking water bath. After collagenase digestion, cell pellets were washed once with “wash buffer” consisting of Ham’s F-10 Nutrient mix (Thermo Fisher Scientific, 22390025) supplemented with 10% Horse serum (Thermo Fisher Scientific, 16050122). This was followed by Dispase digestion with F10 containing 0.4% Dispase II (Thermo Fisher Scientific, 17105041) and 0.2% Collagenase type 2 for 30 mins in a 37°C shaking water bath. After 30 mins incubation, remaining cells were filtered and washed several times. The final cell pellet was re-suspended in PBS containing 2% FBS and satellite cells carrying *Pax7-nGFP* reporter were purified using an SH800S FACS-Sorter (Sony Biotechnologies).

### Myoblast culture

Myoblasts were seeded onto Matrigel-coated plates and cultured in ‘Myoblast medium’. To prepare Matrigel-coated plates, 4% of Matrigel (Corning, 356237) diluted in low glucose DMEM (Thermo Fisher Scientific, 31885023) was applied to the plate which was placed on ice. After 7 mins of incubation on ice, Matrigel was removed from the plate which was further incubated for 1 hr at 37°C. For ‘Myoblast medium’, high glucose DMEM and F-10 were mixed at a similar ratio and supplemented with 20% FBS, 10% horse serum, 1% Pen-Strep and 10 ng/ml basic FGF as previously described^56^. Only P2-P4 myoblasts were used for the reported analyses.

### Quantitative Real time-PCR (qRT-PCR)

Total RNA was extracted using RNeasy mini kit (Qiagen, 74104). The RNA was subjected to DNase treatment (Qiagen, 79254) and its concentration was measured with a Tecan plate reader. cDNAs were synthesized using a High-Capacity cDNA Reverse Transcription Kit (Thermo Fisher Scientific, 4368814) according to the manufacturer’s protocol. qRT-PCR was performed in a 10 μL reaction containing 10 ng of cDNA, 5 μL of Applied Biosystems PowerUp SYBR Green Master Mix (Thermo Fisher Scientific, A25741) and 0.4 μL of each 10 μM forward and reverse primer for the target genes. *Pgk* was used as a house-keeping gene. The sequence of the primers for each target gene is described in Table S1.

**Table S1.**
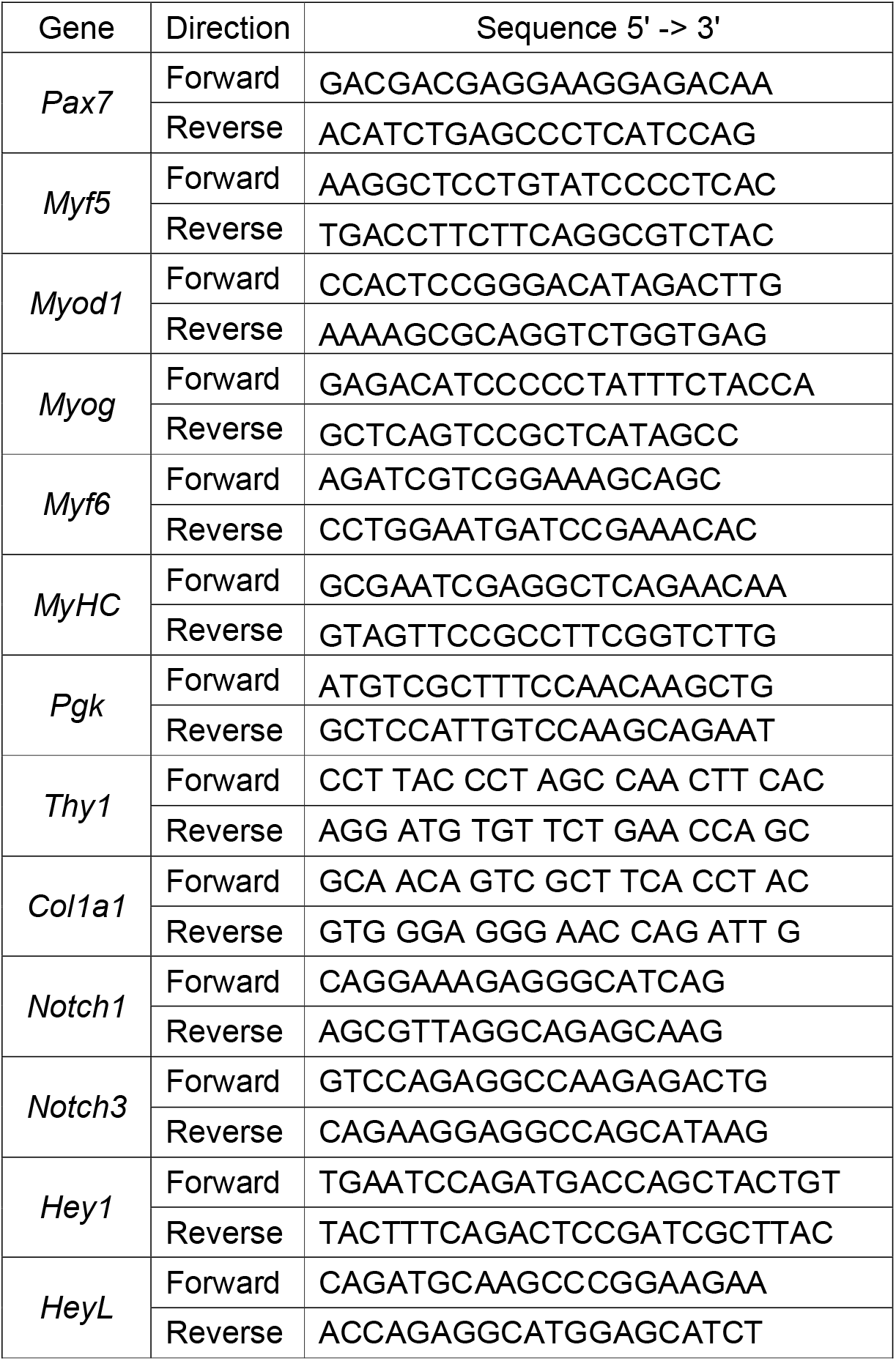
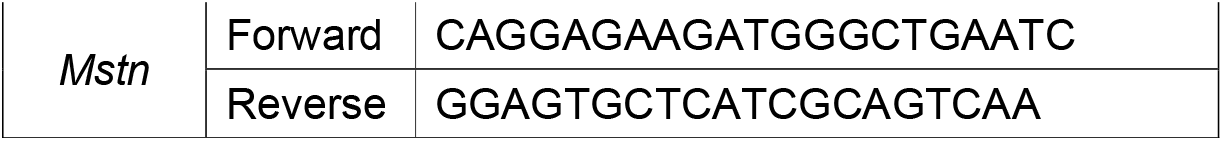
Sequences of the primers used for qRT-PCR

### Immunofluorescence (IF) staining and imaging

Cells were fixed by 4% paraformaldehyde (PFA) (Alfa Aesar, 43368) for 10 mins and incubated in blocking PBS solution containing 2% BSA (AppliChem, 9048-46-8) and 0.2% Triton™ X-100 (Sigma-Aldrich, 9002-93-1) for 30 mins to permeabilize cell membrane and block unspecific antigen binding. As next step, the cells were incubated with primary antibodies in blocking solution for 2 hrs, followed by 30 mins incubation with secondary antibodies and DAPI for nuclei staining (1:1000, Thermo Fisher Scientific, 62248). Stained cells were imaged using a Nikon microscope (ECLIPSE Ti2). The following primary antibodies have been used in this study: Anti-Human/Mouse/Rat/Chicken Pax7 (5 μg/ml, R&D Systems, MAB1675), Anti-Human/Mouse Myod1 (5.8A) (1:100, Thermo Fisher Scientific, MA512902), Anti-Mouse Myosin Heavy Chain (1:1000, R&D Systems, MAB4470) and Anti-Ki-67 (SP6) (1:250, Thermo Fisher Scientific, MA514520). The following secondary antibodies were used in this study at 1:500 dilution: Anti-Mouse IgG1 (Goat, Alexa Fluor 647) (Thermo Fisher Scientific, A21240), Anti-Mouse IgG2B (Goat, Alexa Fluor 546) (Thermo Fisher Scientific, A21240) and Anti-Rabbit IgG (H+L) (Donkey, Alexa Fluor 546) (Thermo Fisher Scientific, A10040).

### Immunofluorescence image quantification

The quantification of immunofluorescent images was performed by counting positive and negative cells. The percentage of the positive cells for each protein was calculated by normalizing the number of positive cells by that of DAPI positive cells.

### Inhibition of Notch via DAPT treatment during reprogramming

For assessment of DAPT treatment during MEF reprogramming, 2.5×10^5^ - 3.0×10^5^ of Rep-MEFs were seeded on 6 well plates and exposed to MyoD or MyoD+F/R/C conditions. For the MyoD+F/R/C+DAPT condition, DAPT (Sigma-Aldrich, D5942) was additionally added to the MyoD+F/R/C condition at a final concentration of 15 μM. All analyses associated with MEF reprogramming in the presence of DAPT were performed at day 10 of treatment. For iMPC cultures, identical number of iMPCs were seeded onto 24 wells plate and cultured in ‘iMPC medium’ either without DAPT (-DAPT) or with DAPT (+DAPT) at the same concentration and conducted 5 days after DAPT treatment began.

### Bulk RNA-Sequencing (RNA-Seq)

Total RNA was extracted using RNeasy mini kit according to the manufacturer’s protocol. The RNA quality was determined by measuring the 28s/18s ratio with a Fragment Analyzer (Agilent, Santa Clara, California, USA). Samples with a 28S/18S ratio over 1.9 were used for library preparation. Libraries were constructed according to the TruSeq Stranded mRNA protocol (Illumina, Inc., California, USA). Briefly, 100-1000 ng of total RNA was poly-A enriched and reverse-transcribed into double-stranded cDNA. Then, the cDNA was fragmented, end-repaired and adenylated before ligation of TruSeq adapters containing unique dual indices (UDI) for multiplexing. Fragments containing TruSeq adapters on both ends were selectively enriched with PCR. The quality and quantity of the enriched libraries were validated using a Fragment Analyzer, which resulted in an average fragment size of approximately 360 bp. The libraries were normalized to 10 nM in Tris-Cl buffer (10 mM, pH8.5) with 0.1% Tween 20 and sequenced with 100bp single end reads on Novaseq 6000 (Illumina, Inc., California, USA) according to standard protocol. Around 20 million reads were obtained for each sample.

### Bulk RNA-Seq data analysis

Raw sequencing reads were pre-processed using the SUSHI framework which was developed in the Functional Genomics Center Zurich^57^ The reads were trimmed (adapter sequences, low quality end and low-quality reads with phred score < 20) first using Trimmomatic v0.36^58^. Pseudoalignment of the trimmed reads was performed against the reference mouse genome assembly GRCm38.p6 and gene expression level (GENCODE release 23) was quantified using Kallisto v0.44^59^. Differential gene expression analysis was conducted between different time points as well as different conditions based on negative binomial distribution using the R package edgeR v3.28^60^. Genes showing variable expression with adjusted (Benjamini-Hochberg method) p-value < 0.05 and two-fold change were considered to be differentially expressed. Gene expression time series data were subjected to soft clustering^61^ using the R package Mfuzz v2.50^62^ to reveal the underlying hidden expression patterns in MyoD and MyoD+F/R/C conditions. For each fuzzy cluster, genes were annotated by gene ontology (GO) terms using the STRING v11^63^ database.

### Protein extraction and digestion

Cultured cells were collected using a cell scraper, snap-frozen in dry ice for 30 mins and stored at −80°C. For protein extraction and digestion, cell pellets were lysed by using a commercial iST Kit (PreOmics, Germany) according to the updated version of the manufacturer’s protocol. Briefly, 100 - 200 μg of the cell pellets were solubilized in ‘Lyse’ buffer, boiled at 95°C for 10 mins and processed with High Intensity Focused Ultrasound (HIFU) for 30s under the ultrasonic amplitude to 85%. Then cell lysates were transferred to the cartridge and digested by adding 50 μL of the ‘Digest’ solution. After an hour of incubation at 37°C, 100 μL of ‘Stop’ solution was added to halt protein digestion. The cartridge was centrifuge at 3800 xg and the through-put was discarded. The peptides remaining in the iST-filter of the cartridge were washed, eluted, dried and re-solubilized in 20 μL of ‘LC-Load’ buffer for MS-Analysis.

### Liquid chromatography-mass spectrometry (LC-MS) analysis

Mass spectrometry analysis was performed on an Orbitrap Fusion Lumos (Thermo Fisher Scientific) equipped with a Digital PicoView source (New Objective) and coupled to a M-Class UPLC (Waters). Channel A was filled with 0.1% formic acid in water while channel B filled with 0.1% formic acid and 99.9% acetonitrile. For each sample, 1-2 μL of peptides were loaded on a commercial MZ Symmetry C18 Trap Column (100Å, 5 μm, 180 μm x 20 mm, Waters) followed by nanoEase MZ C18 HSS T3 Column (100Å, 1.8 μm, 75 μm x 250 mm, Waters). After 3 mins of initial hold at 5% B, a gradient from 5 to 22% B in 83 mins and 22 to 32% B in additional 10 mins was applied. The column was cleaned after the run by increasing to 95% B and holding 95% B for 10 mins prior to re-establishing loading condition. Samples were acquired in a randomized order. The mass spectrometer was operated in data-dependent mode (DDA) acquiring a full-scan MS spectrum (300−1,500 m/z) at a resolution of 120,000 at 200 m/z after accumulation to a target value of 500,000. Data-dependent MS/MS were recorded in the linear ion trap using quadrupole isolation with a window of 0.8 Da and HCD fragmentation with 35% fragmentation energy. The ion trap was operated in rapid scan mode with a target value of 10,000 and a maximum injection time of 50 ms. Only precursors with intensity above 5,000 were selected for MS/MS and the maximum cycle time was set to 3 s. In charge state screening, singly assigned and unassigned charge states, and charge states higher than seven were rejected. Precursor masses previously selected for MS/MS measurement were excluded from further selection for 20 s, and the exclusion window was set at 10 ppm. The samples were acquired using internal lock mass calibration on m/z 371.1012 and 445.1200. The data obtained from the mass spectrometry proteomics were handled using the local laboratory information management system (LIMS)^64^.

### Protein identification and label free protein quantification (LFQ)

The acquired raw MS data were processed by MaxQuant (version 1.6.2.3), followed by protein identification using the integrated Andromeda search engine^65^. Spectra were searched against a Swissprot Mus musculus reference proteome (taxonomy 10090, version from 2019-07-09), concatenated to its reversed decoyed fasta database and common protein contaminants. Carbamidomethylation of cysteine was treated as a fixed modification, while methionine oxidation and N-terminal protein acetylation treated as a variable. Trypsin/P was set for enzyme specificity, allowing a minimal peptide length of 7 amino acids and a maximum of two missed-cleavages. MaxQuant Orbitrap default search settings were used. Peptides with FDR < 0.01 and proteins with FDR < 0.05 were processed for further steps. Label free quantification was carried out by applying a 2 minutes window for match between runs. LFQ intensity results from MaxQuant were used for a hierarchical clustering across all samples implemented in Perseus software^66^.

### Two-group comparison analysis in proteomics

Each sample file was kept separate in the experimental design to acquire individual quantitative values in the MaxQuant experimental design template. Fold changes of proteins were calculated based on intensity values reported in the proteinGroups.txt file. Filtration for proteins with 2 or more peptides allowing a maximum of 4 missing values, normalization with a modified robust z-score transformation and the t-test with pooled variance to compute p-values were implemented using the functions in R package SRMService^67^. If all measurements of a protein are missing in one of the conditions, a pseudo fold change was calculated replacing the missing group average by the mean of 10% smallest protein intensities in that condition. Proteins showing variable expression with adjusted p-value < 0.05 and two-fold change were considered to be differentially expressed between the conditions.

### Bulk RNA-seq / proteomics correlation analysis

An integrated dataset was created with differentially expressed genes overlapping with differentially expressed proteins based on their common Ensembl identifiers. Pearson correlation coefficient was computed between differentially expressed genes and proteins using their log2 fold change values.

### Enrichment analysis for functional annotation

For bulk RNA-seq data, the pathway enrichment analysis was carried out using Gene Set Enrichment analysis (GSEA) based on genes ranked in order of log2FC and the functional WikiPathways (www.wikipathways.org) database via WEB-based GEne SeT AnaLysis Toolkit (WebGestalt)^68^. Only significant (FDR < 0.05) pathway categories with ten to 500 genes were considered for enrichment analyses. For proteomics and integration between bulk RNA-Seq and proteomics datasets, enrichment analysis was performed based on significant DEGs / DEPs (|Log2FC|>1, adj.P-value < 0.05) in Metacore (https://clarivate.com/products/metacore/, Clarivate Analytics, London, UK). Only significant (FDR < 0.05) pre-built process networks were presented. For bulk RNA-seq data comparison between Pax7-nGFP^+^ iMPCs and Pax7-nGFP^+^ myoblasts, overrepresentation analysis (ORA) was performed using the R package goseq v1.42^69^.

### ATAC-Seq

Libraries for ATAC-Seq were constructed using Omni-ATAC protocol as previously reported^70^. Briefly, 50,000 – 60,000 cells were lysed on ice by cold lysis buffer containing 0.1% NP-40 (Sigma-Aldrich, 98379), 0.1% Tween-20 (Sigma-Aldrich, P3416), 0.01% Digitonin (Promega, G9441), 10 mM Tris-HCl (pH 7.5), 10 mM NaCl, 3 mM MgCl_2_ in Nuclease-free water. Nuclei were collected by centrifugation and subjected then to transposition reaction in 1x Tagment DNA buffer containing 5% Tn5 Transposase (Illumina, 20034197), 0.1% Tween-20, 0.01% Digitonin at 37°C for 30 mins on thermomixer, followed by DNA purification using MinElute PCR purification kit (Qiagen, 28004). To identify samples, index PCR amplification was performed by mixing 10 μL of transposed DNA with 10 μL nuclease-free water, 25 μL NEBNext High-Fidelity 2X PCR Master Mix, 2.5uL of each 25 μM Ad.1 and Ad.2, as previously published^71^. Total PCR cycle was determined according to the amplification graph after the first 12 cycles. To remove primer dimers and larger fragment, double-sided bead purification was carried out using AMPure XP (Beckman Coulter, A63881). Library quality was validated using TapeStation (Agilent Technologies) and sequenced on HiSeq-2500 (Illumina, Inc, California, USA) with paired end of 70bp. All libraries had more than 50 million reads.

### ATAC-seq analysis

ATAC sequencing was performed on a HiSeq2500 instrument at FGCZ. After initial quality control (adapter and low-quality base trimming) using fastp v0.20^72^, raw sequencing reads were mapped against the reference mouse genome assembly GRCm38.p6 using Bowtie2 v2.4.1^73^. PCR duplicates were removed using the MarkDuplicates tool from Picard (https://broadinstitute.github.io/picard/). Peak calling was performed using MACS2^74^ (3) with-nomodel −f BAMPE −gsize mm −keep-dup all −extsize 200 options. Called peaks were annotated using the R package ChIPseeker^75^. For each sample, fragment count matrix was generated using the R package chromVAR^76^ based on the promoter region, defined as −1 to 1 kb around the transcription start site. The R package edgeR v3.28^60^ was used to perform differential expression analysis between different conditions using the fragment count matrices. Genes showing variable expression with p-value < 0.01 and one-fold change were considered to have an open chromatin in the defined promoter region.

### Bulk RNA / ATAC-seq correlation analysis

An integrated data set was created with differentially expressed genes overlapping with the genes with open chromatins in the promoter region based on their common Ensembl identifiers. Pearson correlation coefficient was calculated for the overlapping genes based on their log2 fold change values.

### Single cell RNA-Sequencing (scRNA-Seq)

An early passage iMPC clone was trypsinized and filtered using a 40 μm cell strainer (VWR, 734-0002) to filter out debris and fragments of myofibers. Filtered iMPCs were washed with PBS and the number of cells was counted manually using a hemocytometer with Trypan blue (Sigma-Aldrich, T8154) staining. Next, the cell pellet was re-suspended in PBS at a concentration of 1000 cells/μL and immediately used for 10x single cell library construction. 10x library was built using Single cell 3’ reagent kit v3 (10xGenomics, Pleasanton, CA) according to the manufacturer’s protocol. Briefly, cells were loaded in chromium chip B targeting ~10,000 recovered cells. Generated GEMs were cleaned, and cDNA was amplified by PCR, followed by cDNA fragmentation, end repair, A-tailing, adaptor ligation, index PCR, and double sized selection. The library was sequenced on a full SP flowcell of Novaseq 6000 (Illumina, Inc, California, USA) which allows to obtain 560 million reads for around 10,000 cells (over 40,000 reads per cell).

### scRNA-Seq data analysis

CellRanger v4.0.0 pipeline was used for demultiplexing the sample, mapping raw reads to the mouse reference genome (build GRCm38.p6) and generating feature-barcode count matrix^77^. The count matrix was further analyzed using Seurat v3.2.3 pipeline^26,78^. Droplets with unique feature counts < 250 and > 7,200 and mitochondrial gene counts > 5% were discarded for quality control. The filtered data was globally scaled via log normalization. The scaled data was dimensionally reduced via principal component analysis (PCA) using 2,000 highly variable genes. Cells were clustered based on first 20 principal components (PCs) using the Louvain algorithm^79^ with a resolution of 0.5. Uniform manifold approximation and projection (UMAP)^80^ method was applied using the same PCs to visualize the clustered cells in low-dimensional place. Cluster biomarkers were identified using the Wilcoxon rank-sum test (adjusted p-value < 0.05).

### Bulk RNA-Seq / scRNA-Seq correlation analysis

To generate pseudo bulk data from scRNA-seq at the sample level, raw counts were aggregated across clusters of similar cell types (fibroblasts, progenitors). Pearson correlation coefficient was computed between the overlapping genes expressed in bulk and pseudo bulk RNA-seq data based on their log2 normalized expression values.

### Single cell trajectory analysis of scRNA-Seq dataset

Unsupervised single cell trajectory analysis was performed using Monocle 3^81,82^. The cells were ordered in pseudotime along the learned trajectory with cluster 1 being the root. Due to the presence of a strong feature outside the focus of interest which might influence unsupervised analysis, we performed semi-supervised single cell trajectory analysis using Monocle 2^27,81^. Six genes were defined as markers namely *Ccnb1* for cycling progenitors, *Myog* for committed progenitors, *Myog* and *Tnnt2* for myocytes, *Pdgfrb* and *Fbln2* for connective tissue / fibroblast cells and *Pax7* for Pax7^+^ progenitors. A set of differential genes was selected, which co-varies with these markers. Cells were ordered using the top 1,000 differentially expressed genes based on their adjusted p-values. Each cell was assigned a pseudotime value to capture its progress during the biological process. Branch expression analysis modeling^83^ was performed to find the branch-dependent genes that could identify the mechanism underlying the cell fate decisions. Branch-dependent genes categorized by each cluster were annotated with GO term using the online tool DAVID v.6.8^84,85^

### Statistical analysis

We utilized 3 cell lines per group that were isolated from different mice for bulk RNA-Seq and other experiments including iMPC generation and Notch inhibition, whereas 4 cell lines for LC-MS were used. We utilized the software Prism to perform statistical analysis for gene expression results such as qRT-PCR data as indicated in the figure legends. Statistical analysis for multi-omics datasets was carried out as described in each respective section.

## Notes

### Competing Interest Statement

The authors have declared no competing interest.

