## Supplementary materials for "Integrative molecular roadmap for direct conversion of fibroblasts into myocytes and myogenic progenitor cells"

**Integrative molecular roadmap for reprogramming fibroblasts into induced myogenic stem and progenitor cells**

**This file includes:**

Supplementary Figs. 1 to 7

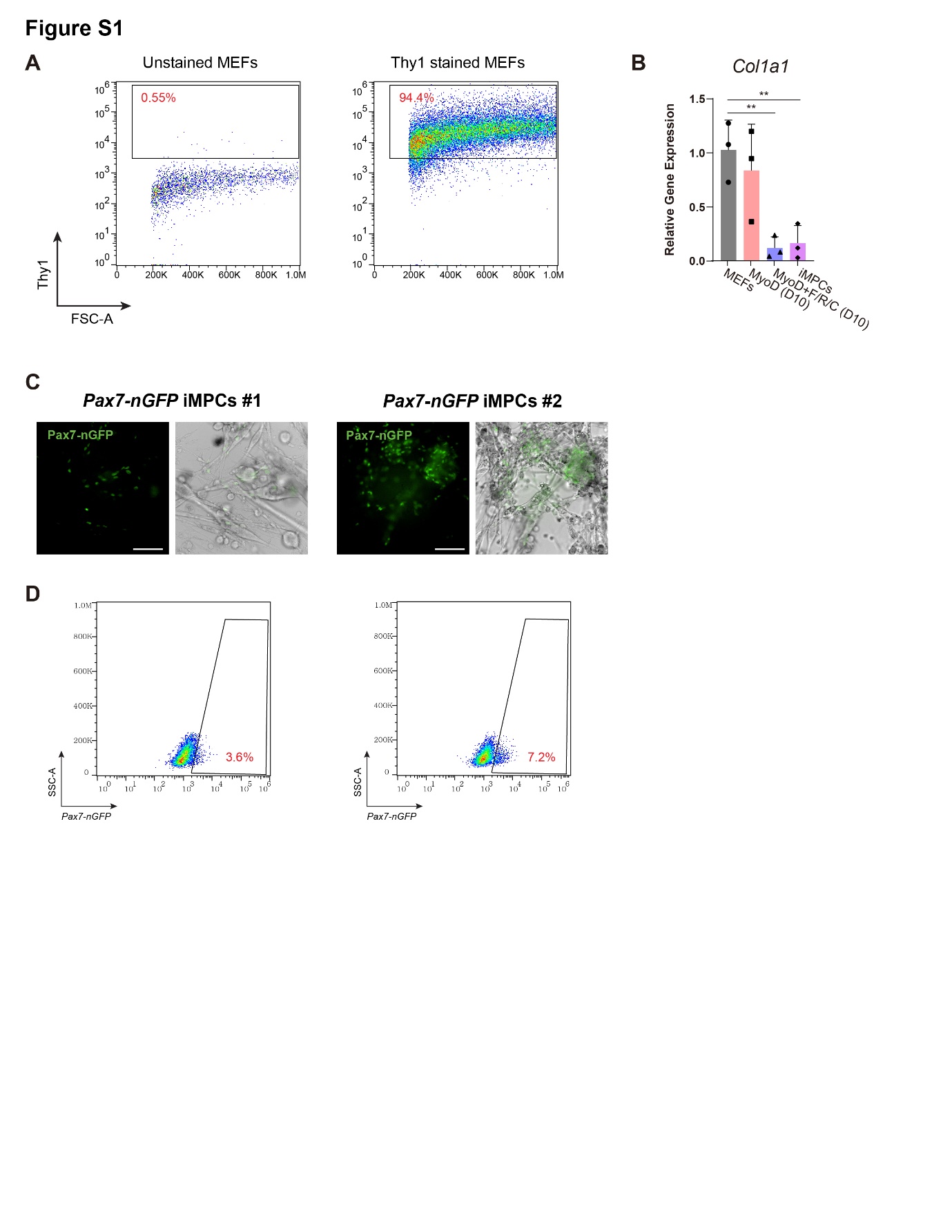

**Supplementary Fig. 1: Reprogramming MEFs into iMPCs via MyoD and small molecule treatment**

**(A)** Flow cytometry analysis of MEF cultures stained for the fibroblast-specific surface marker Thy1. MEFs, mouse embryonic fibroblasts. **(B)** qRT-PCR analysis for the fibroblast-specific marker *Col1a1*. Data is shown as means ± S.D. N = 3 cell lines per each group. Statistical significance was determined by a two-tailed unpaired *t*-test (**p<0.01, ***p<0.001, n.s=non-significant). **(C)** Representative images of two stable *Pax7-nGFP* iMPC clones at passage 1. Note nuclear GFP expression only in mononucleated cells. Scale bar, 100 µm. **(D)** Flow cytometry analysis of two *Pax7-nGFP* iMPC clones containing 3.61% or 7.23% Pax7 positive cells, respectively.

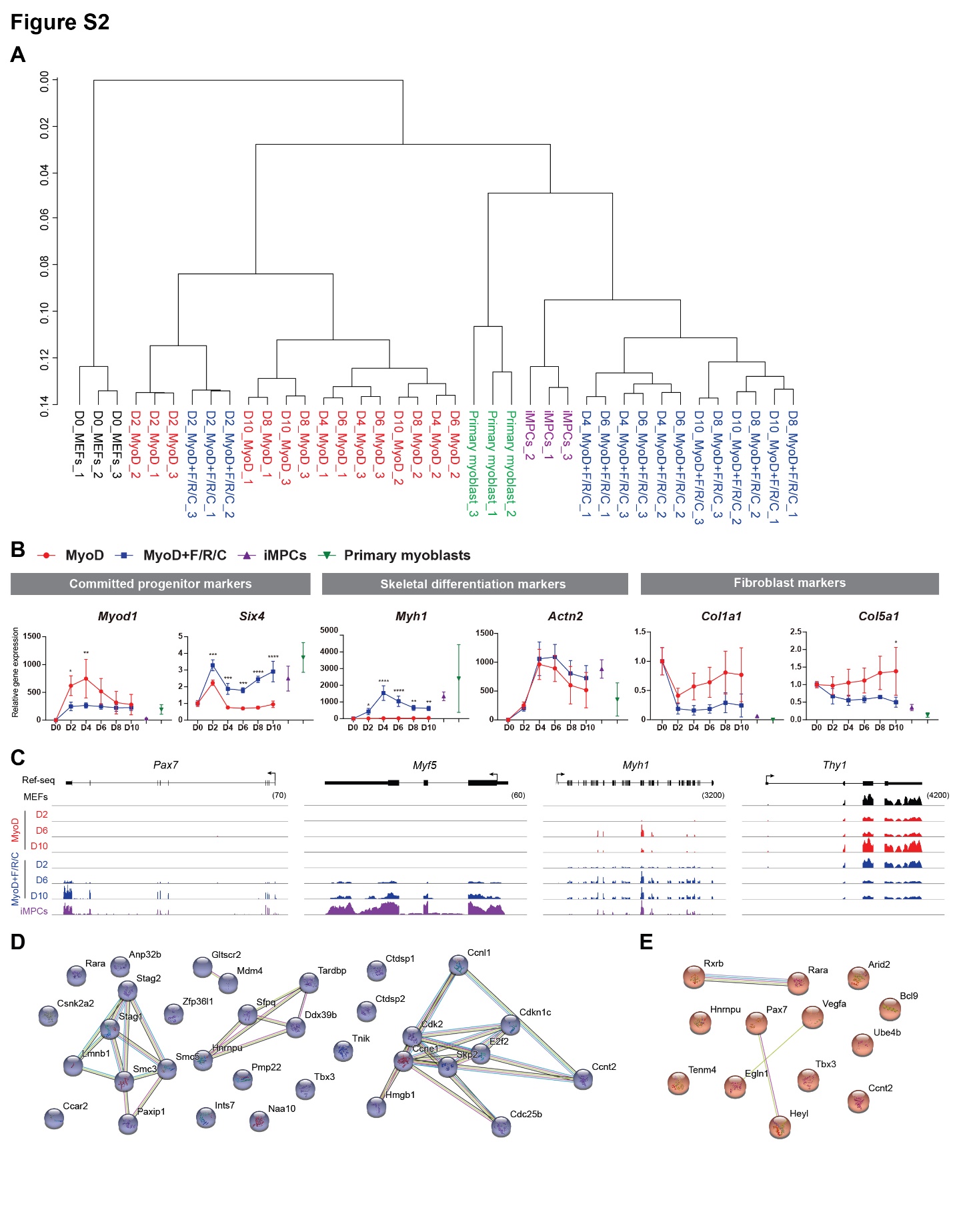

**Supplementary Fig. 2: Gene expression dynamics during transdifferentiation and reprogramming**

**(A)** Dendrogram clustering based on bulk RNA-seq data using all gene read counts for the indicated samples. N = 3 cell lines per each group. **(B)** Gene expression analysis based on bulk RNA-Seq data for the selected genes in MEFs subjected to MyoD or MyoD+F/R/C treatment. Established iMPCs and primary myoblasts served as positive controls. Relative gene expression was calculated by normalizing the RPKM values of each sample to that of parental MEFs (D0 MEFs). The data is shown as means ± S.D. N = 3 cell lines per each group. Statistical significance was determined by two-way ANOVA between conditions at each time point (*p<0.05, **p<0.01, ***p<0.001, ****p<0.0001). **(C)** Representative Integrative Genomic Viewer (IGV) tracks for the indicated genes during a reprogramming time course at the respective time points. **(D-E)** Enriched network analysis using the STRING database for the genes shown in gene set 2 (MyoD+F/R/C treatment, Fig. 2E) that annotated with *Regulation of cell cycle* (D) and *Striated muscle tissue development* (E).

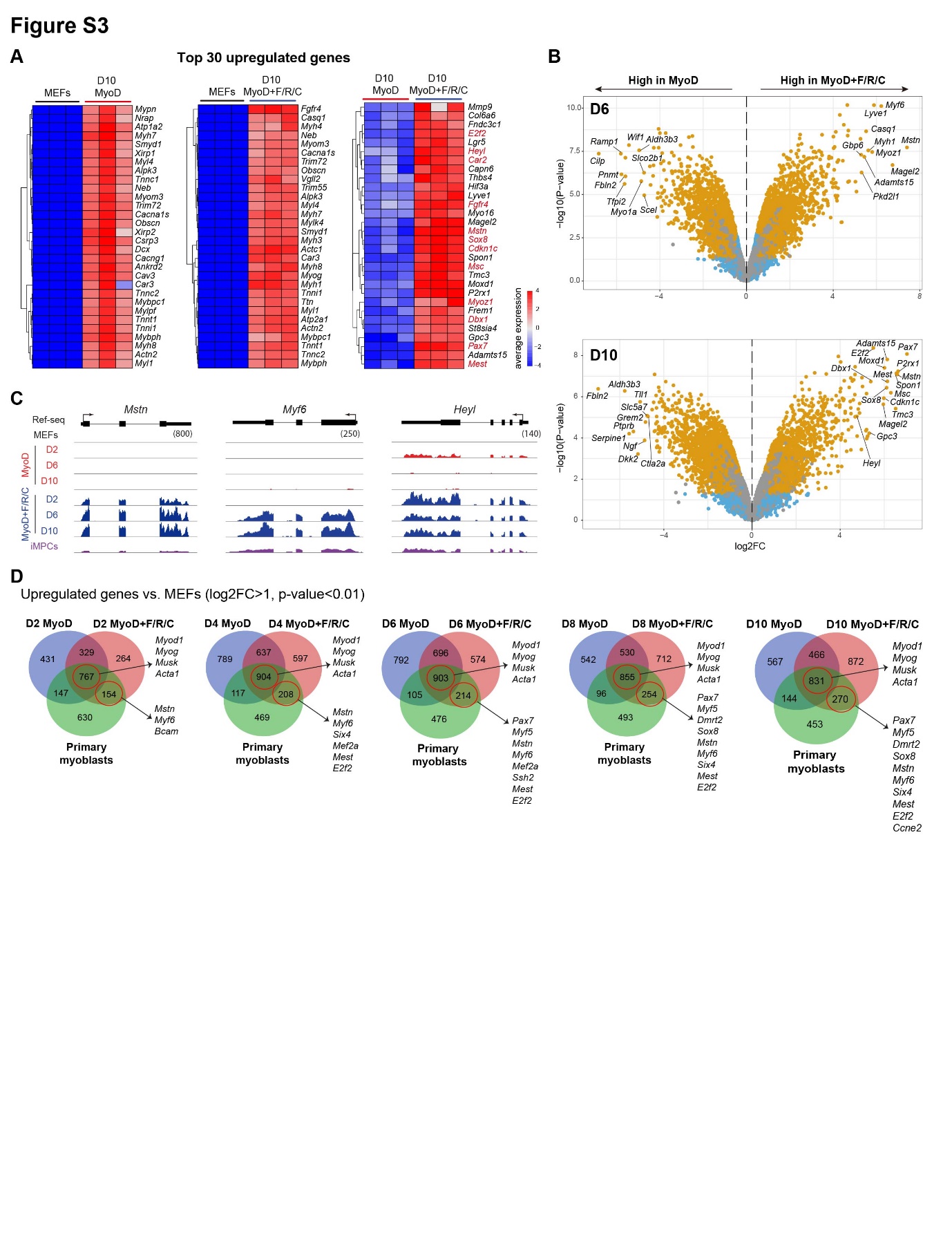

**Supplementary Fig. 3: Transcriptional differences between transdifferentiation and reprogramming**

**(A)** Heatmaps based on bulk RNA-Seq data displaying the top 30 upregulated genes between the indicated cell lines. The average expression is presented as gradient. N = 3 cell lines per group. Unique myogenic genes are highlighted as red color. **(B)** Volcano plots based on bulk RNA-Seq data for differentially expressed genes (DEGs) between MyoD and MyoD+F/R/C treatment at day 6 and 10 of the reprogramming process. Significant DEGs (|log2FC|>0.5, p-value<0.05) are shown as yellow dots. N = 3 cell lines per each group. **(C)** IGV tracks for select genes shown in (B). **(D)** A Venn diagram based on bulk RNA-Seq data showing the overlap of upregulated genes in primary myoblasts and MEFs subjected to MyoD or MyoD+F/R/C conditions at each time point vs. parental MEFs (log2FC>1, p-value<0.01). N = 3 cell lines per each group.

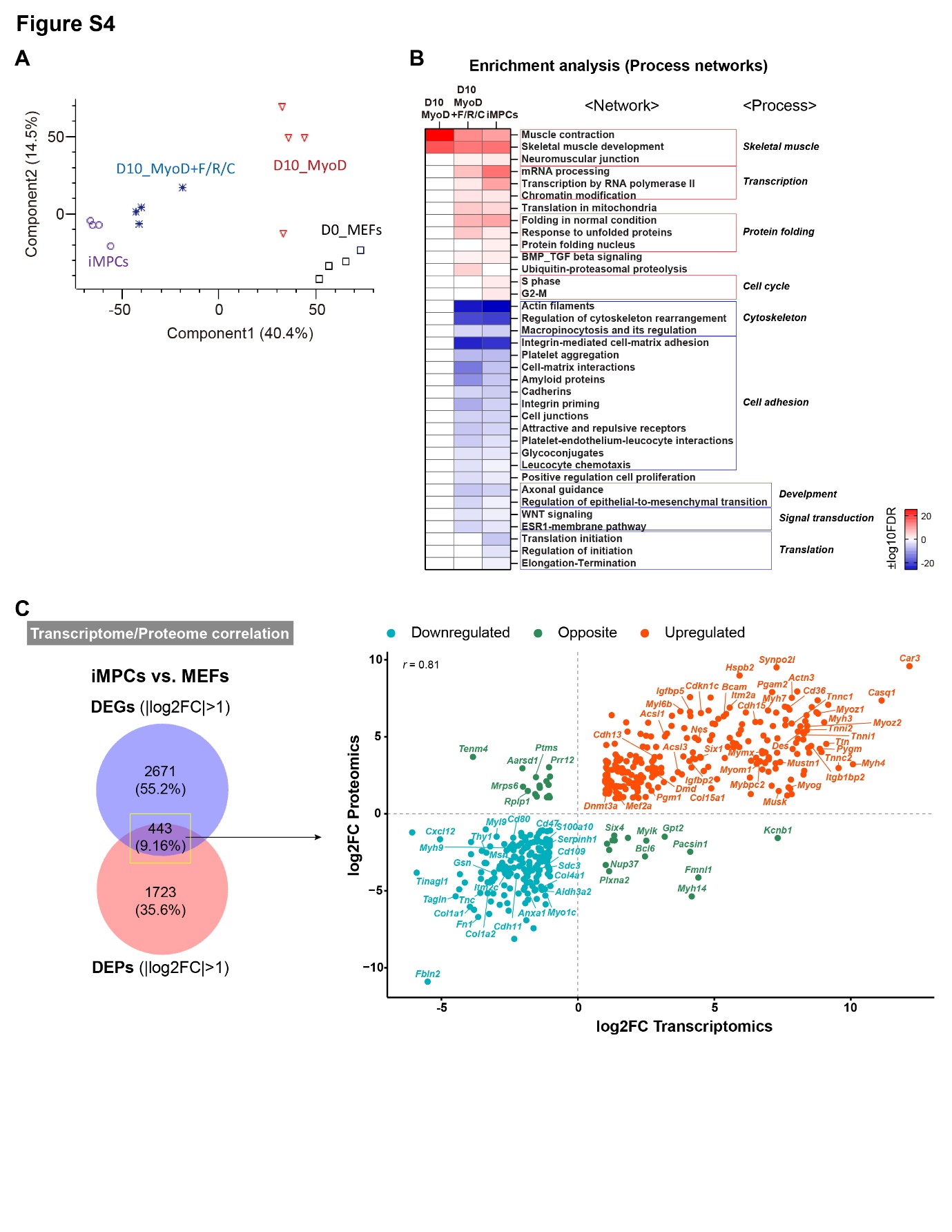

**Supplementary Fig. 4: Proteomic analysis during transdifferentiation and reprogramming**

**(A)** PCA of global protein expression based on LC-MS analysis across all samples. N = 4 cell lines per each group. **(B)** Enriched process networks of DEPs in each condition vs. MEFs (|log2FC|>1, p-value<0.05) as analyzed in Metacore. Only significantly enriched Process networks are presented (FDR<0.05). Upregulated (Red) and downregulated (Blue) process networks are shown using -log10(FDR) and log10(FDR), respectively. **(C)** Left- A Venn diagram based on transcriptome and proteome datasets showing the number of preselected and significant DEGs and DEPs in iMPCs vs. MEFs (|log2FC|>1, p-value<0.05). Right- scatterplot showing the correlation between the transcriptome and proteome datasets of iMPCs vs. MEFs. In total, 443 overlapped DEGs / DEPs are projected on the plot, corresponding to the Venn diagram analysis shown on the left. Red and blue dots indicate upregulated and downregulated genes or proteins, respectively.

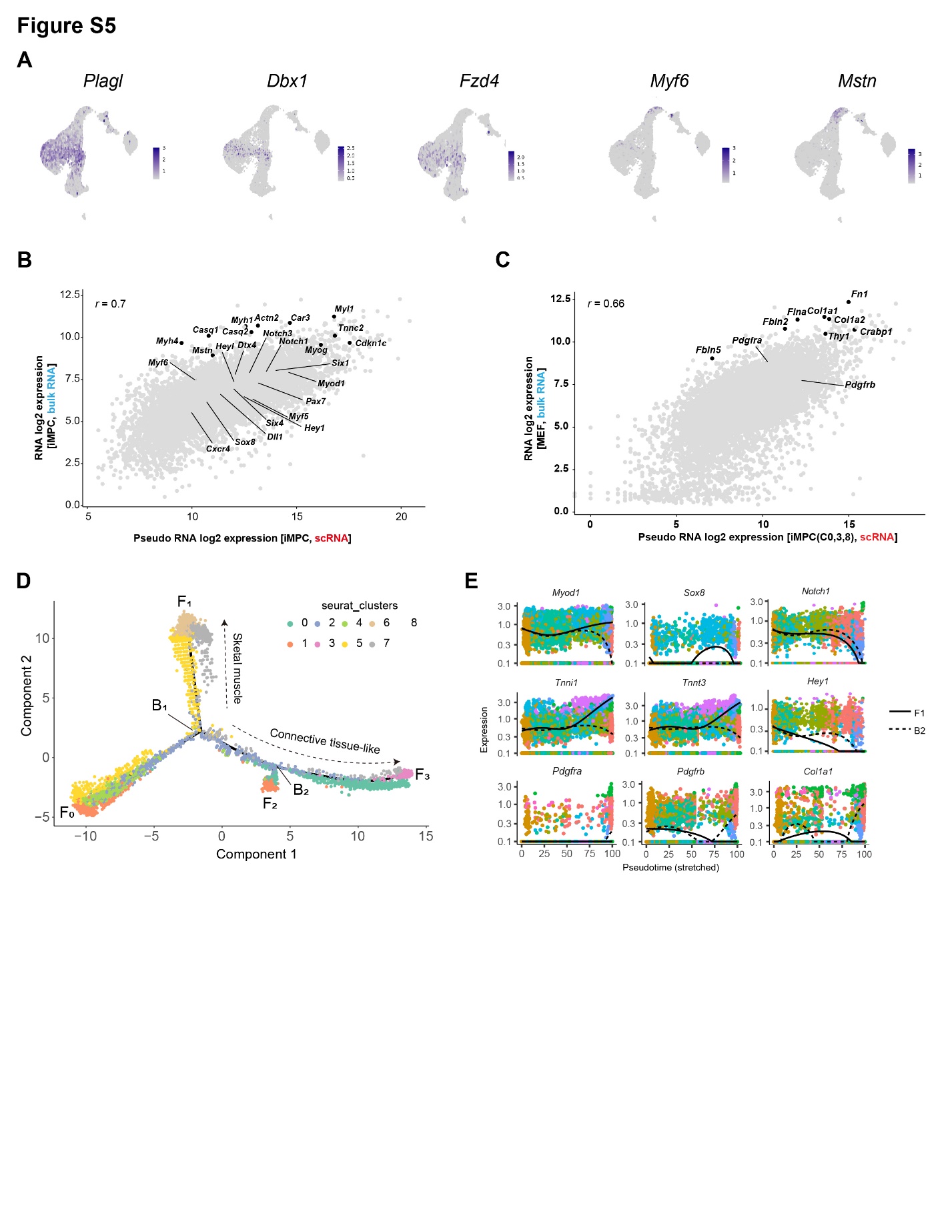

**Supplementary Fig. 5: scRNA-Seq analysis of iMPCs**

**(A)** UMAP plot for the indicated genes based on scRNA-Seq data. **(B-C)** Scatter plot displaying gene expression correlation profile between scRNA-Seq and bulk mRNA data for an iMPC clone (B) and MEFs (C). Pseudo bulk mRNA expression was computed from scRNA-Seq data for the indicated clusters. The cluster identities correspond to the UMAP projection shown in Fig. 5A. **(D)** Minimum spanning trees showing ordered cells based on semi-supervised single cell trajectory analysis reconstructed by Monocle2 and colored by cell cluster identifiers. **(E)** Plot showing the expression of the indicated genes as a function of pseudotime. Dots indicate cells colored by cell identifiers.

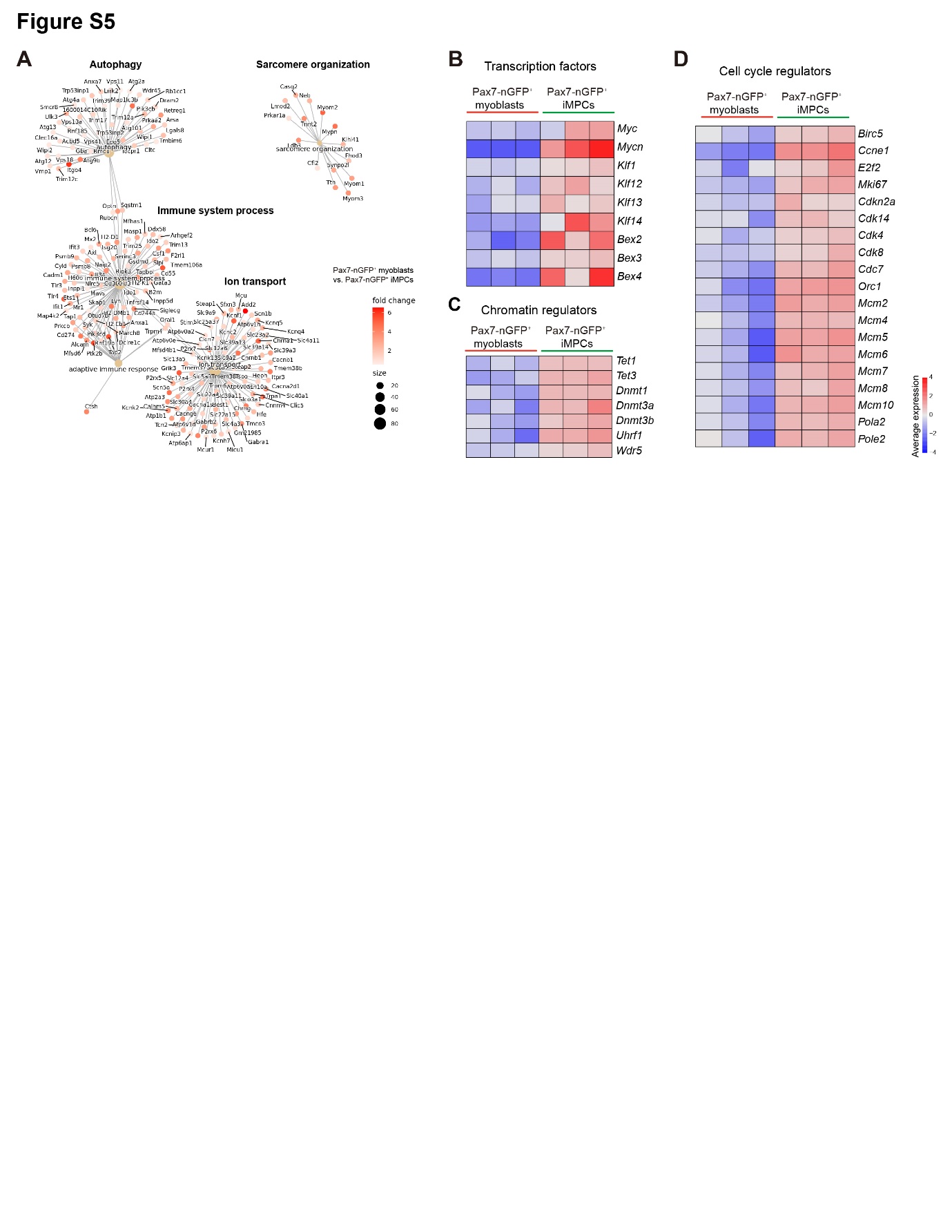

**Supplementary Fig. 6: Molecular analysis of Pax7^+^ iMPCs and Pax7^+^ primary myoblasts**

**(A)** Over Representation Analysis (ORA) of the gene sets that were significantly enriched (log2FC > 0.5, p-value < 0.01) in Pax7-nGFP^+^ myoblasts in comparison to Pax7-nGFP*^+^* iMPCs. Only significant GO biological processes (adj. p-value < 0. 05) are shown. The size indicates the number of genes involved in the biological process and the color-coding scale represents the fold change between the two conditions. **(B-D)** Heatmap of selected genes associated with transcription factors (B), chromatin regulators (C) and cell cycle regulators (D) that were highly expressed in Pax7-nGFP*^+^* iMPCs in comparison to Pax7-nGFP*^+^* myoblasts. Gradient of high to low expression of each gene relative to the average expression in each comparison is indicated by red to blue. N = 3 cell lines per each group.

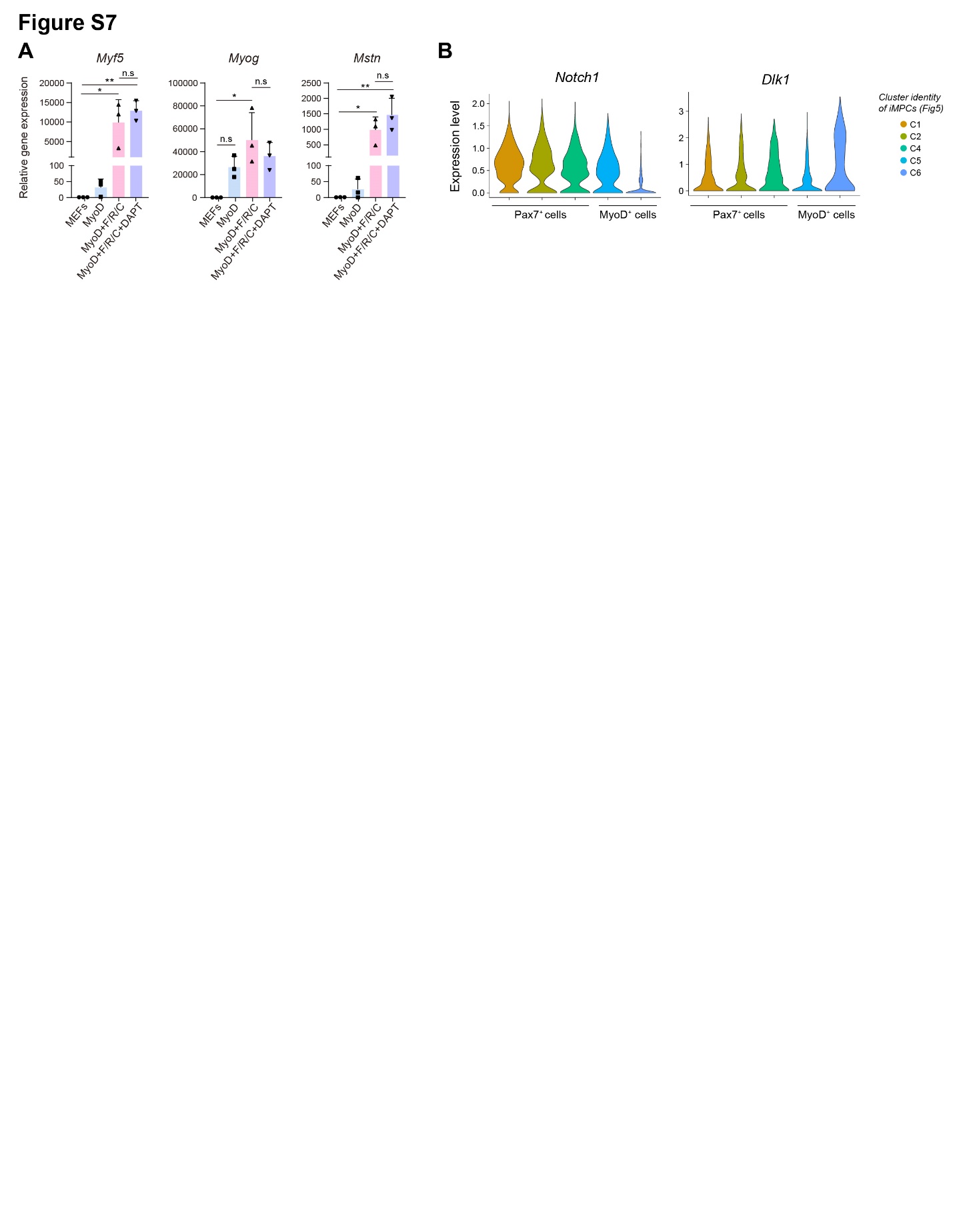

**Supplementary Fig. 7: The Notch pathway is critical for iMPC reprogramming**

**(A)** Relative qRT-PCR gene expression analysis for the indicated myogenic genes at day 10 of MEF reprogramming using the indicated conditions. The data is shown as means ± S.D. N = 3 cell lines per each group. Statistical significance was determined by one-way ANOVA (*p<0.05, **p<0.01, n.s=non-significant). **(B)** Violin plots showing the average expression of *Notch1* and *Dlk1* in the respective clusters based on scRNA-Seq data from Fig 5.
